# Phenol Sensing in Nature Modulated via a Conformational Switch Governed by Dynamic Allostery

**DOI:** 10.1101/2022.05.18.492265

**Authors:** Jayanti Singh, Mohammad Sahil, Shamayeeta Ray, Criss Dcosta, Santosh Panjikar, G. Krishnamoorthy, Jagannath Mondal, Ruchi Anand

**Author notes:** **For correspondence Ruchi Anand**, Department of Chemistry, Indian Institute of Technology Bombay, Mumbai-400076, India;, **Jagannath Mondal**, Centre for Interdisciplinary Science, Tata Institute of Fundamental Research, Hyderabad-500046, India.

## Abstract

NtrC family of proteins sense external stimuli and accordingly stimulate stress and virulence pathways via activation of associated σ54-dependent RNA polymerases. Here, we establish that MopR, an NtrC protein, harbors a dynamic bi-directional electrostatic network that connects the phenol pocket to two distal regions, namely the “G-hinge” and the “allo-steric-linker”. While G-hinge influences the entry of phenol, the allosteric-linker passes the signal to the downstream ATPase domain. Phenol binding induces a rewiring of the electrostatic connections by eliciting dynamic allostery, and it was demonstrated that perturbation of the core relay residues results in a complete loss of ATPase stimulation. A mutation of the G-hinge,∼20Å from the phenol pocket, demonstrated altered flexibility by shifting the pattern of conformational states accessed, leading to a protein with 7-fold enhanced phenol binding ability and enhanced transcriptional activation. A global analysis illustrates that dynamic allostery-driven conserved community networks are universal and evolutionarily conserved across species.

## INTRODUCTION

Proteins are involved in several biological processes where they undergo various conformational fluctuations, modulated by diverse signals such as ligand binding, environmental factors, etc.^1–3^ Protein breathes through complex multi-dimensional landscape, which constitutes various conformational states.^4,5^ For instance, allostery in proteins where binding of a ligand/effector at a distal site can stabilize a particular conformation can be one of the strategies adopted by nature to enable function.^6^ Here, long distance communication can either be governed by conformational allostery which is accompanied by a significant structural change within the protein structure or it can be dynamic where no detectable conformational change is observed.^7–12^. While in the case of conformational allostery, enthalpic contributions play a central role, in dynamic allostery entropy effects are considered to be predominant.^8^ This is due to the fact that energetic perturbations, which are generally caused by small scale internal sidechain changes, can result in reorganization of the interaction network of the whole protein.^7,13^ Most commonly used techniques to understand allosteric mechanism are by combining structure determination tools such as X-ray crystallography, NMR and more recently cryo-electron microscopy with biophysical and computational tools. The structural tools partly provide snapshots of selective conformations of a protein, within its functional cycle, the supporting techniques help in delineating associated conformational heterogeneity that together help in unravelling the mechanism of complex allosteric regulation.

NtrC family of protein belongs to the broader bacterial enhancer binding proteins that assemble into ATPase motors. They serve as mechanoenzymes by triggering activation of σ54-dependent RNA polymerase(RNAP).^14,15^ They respond to external stimuli and via either phosphorylation of their signal sensing domain or by entry of a particular ligand and in some cases via protein-protein interaction activate the RNAP for downstream transcription of select pathways.^15^ MopR, a NtrC family protein from *Acinetobacter calcoaceticus* NCIB8250 is a multi-domain protein where binding of phenol to its N-terminally located ligand binding [A] domain results in de-repression of ATP hydrolysis via the centrally located AAA+[C] domain.^14,16^ It also harbors a C-terminal DNA binding [D] domain which assists MopR to latch onto a specific DNA (Figure 1a, S1a)^14,16^ segment, upstream activation site (UAS), that lies 100-200 bases upstream of the RNAP binding region. It is hypothesized that in presence of phenol and ATP, MopR assembles into an oligomeric structure and executes its mechano-function by activating the σ^54^-RNAP which then triggers the downstream phenol degradation pathway. This ability of MopR to sense phenol and elicit a response has been used to create a plethora of sensors for monitoring concentrations of several xenobiotics in polluted water.^14,17–19^ The ligand binding domain of MopR is connected to the AAA+ domain via a connector B-linker helix and it is envisioned that binding of phenol brings about a global allosteric change, transmitted via the B-linker, that is crucial for activating ATP hydrolysis thereby, allowing MopR to elicit its function.^14,20,21^

**Figure 1.**
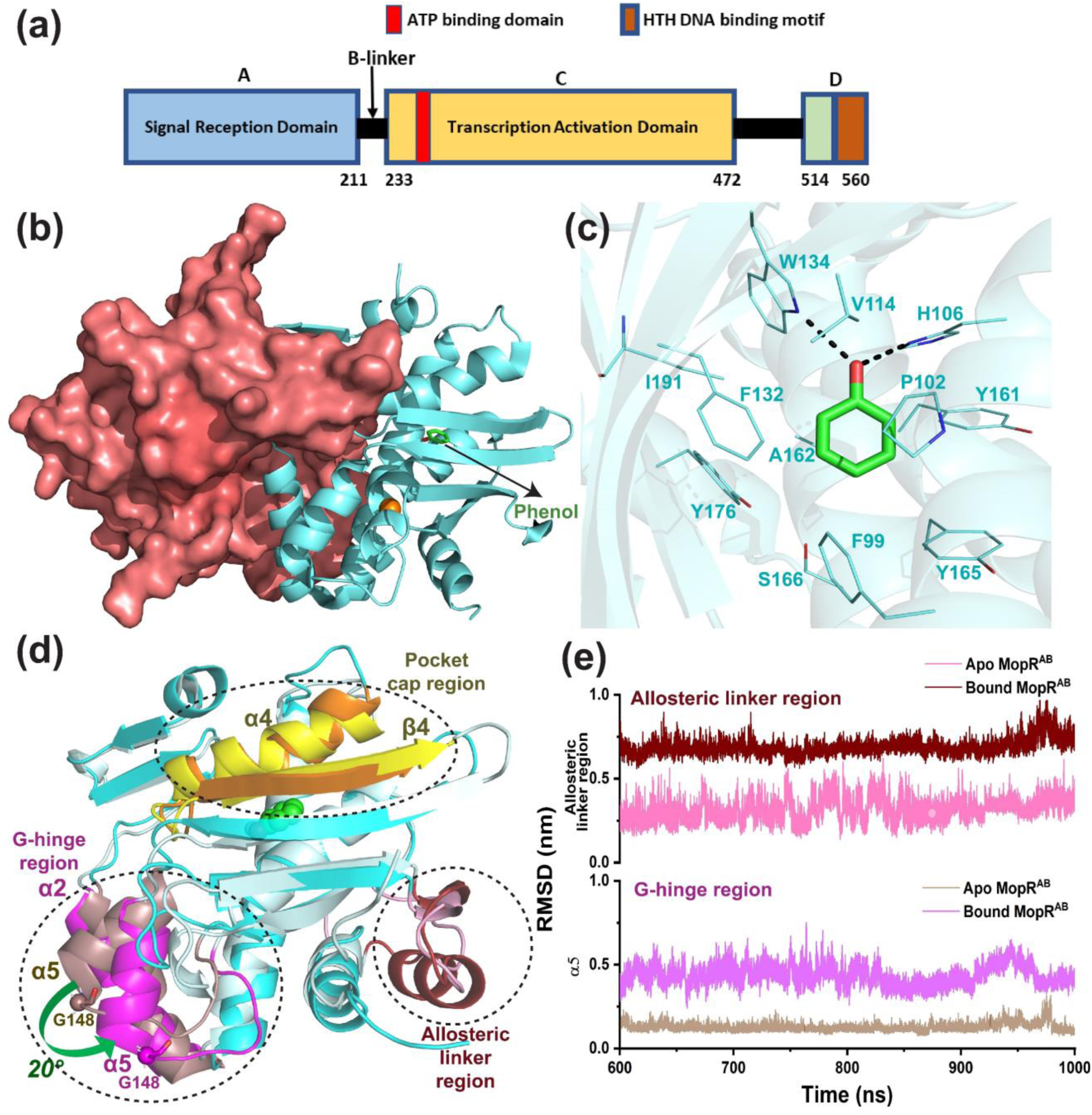
**(a)** Domain organization of MopR. **(b)** MopR^AB^ (residues 1-229) (PDB ID: 5KBE) dimer with one protomer in cartoon representation and other in surface representation (deep salmon) showing phenol completely buried. **(c)** Phenol binding pocket of MopR^AB^. **(d)** Superimposition of a representative snapshots, 900ns and 950ns of the apo (pale blue) and bound (cyan) MopR^AB^ structures. The G-hinge region is shown in pale magenta and magenta for apo and bound MopR^AB^, respectively. The allosteric linker region is shown in light pink and wine red for apo and bound MopR^AB^ **(e)** RMSD plot of apo and bound MopR^AB^ depicting conformational changes in the α5 (G-hinge) and allosteric linker region.

There has been paucity in obtaining structural information in this family of enzymes. The structure of the signal sensing domain [A] and a portion of the tandem B-linker of MopR (MopR^AB^) (Figure S1b) was determined in complex with its cognate ligand, phenol (Figure 1b, PDB ID:5KBE) and a few of its derivatives in 2016.^14^ The structure revealed that the ligand binding domain of MopR belongs to the nitric oxide signaling fold that encompasses closely related bacterial homologs such as dimethylphenol regulator (DmpR) that responds to 2,3-dimethylphenol and XylR that senses benzene. MopR also showed structural similarity to evolutionarily distant eukaryotic proteins which bind fatty acids and are part of a large complex that partakes in transport across the golgi.^14,22^ The [A] domain of MopR harbors a zinc binding site, which is ∼10 Å away from the phenol binding region, whose function is attributed to maintain structural integrity.^14^ The structure was especially instrumental in understanding the key features that lead to selective binding of phenol. In brief, the binding pocket of phenol is mostly composed of hydrophobic residues that stabilizes the aromatic ring of phenol via п-stacking interactions.^14^ In particular the phenolic OH moiety is anchored in the active site via hydrogen-bonding interactions by key sensor residues, H106, and W134 (Figure 1c). The snug nature of the pocket imparts it selectivity such that mono-substituted phenols bind with slight reduction in binding affinity whereas, bulkier phenols are unable to bind.^14,17–19^ The OH anchor ensures that non-phenolic aromatic hydrocarbons are unable to occupy the phenol pocket.^18,19^

More recently the X-ray crystal structure of a homolog of MopR, DmpR, that encompasses both the sensor along with the ATPase domain has also been determined.^23^ In this structure the relative orientations of the [A] and the central AAA+ [C] domains when phenol is bound to the [A] domain could be visualized. However, since most AAA+ motors assemble as hexamers or heptamers it is difficult to discern if the captured tetrameric form is an inactive or an activated conformation of the enzyme.^15,24,25^ The structure of the unliganded form of MopR and its homologs has not been determined till date. Since MopR is very selective towards its ligands, here we explore conformational changes that enable the enzyme to bind its ligand. Moreover, how ligand binding initiates de-repression of the downstream ATPase domain, the associated signal transduction pathway that connects the two domains is also not well understood, resulting in the overall mechanism to remain elusive. Therefore, as a first step to understand how the binding of ligand (phenol), induces significant conformational changes both within the [A] domain of the protein as well as those that percolate outside to the adjacent B-linker region, we carried out a combination of experimental and molecular dynamic (MD) simulation studies. Further, regions of the enzyme that reside outside the phenol pocket that may influence and play a critical role in facilitating function were also explored. The study in particular helped in unravelling how dynamic networks that run within the protein play a critical role in communication. More importantly it establishes a conserved community network relay that also exists in other aromatic ligand binding subclass of NtrC family of proteins. It especially entails how distal regions far from the binding or active sites play a significant role in flow of information and thereby, control overall protein function.

## RESULTS

### Dynamic networks in MopR

Analysis of the crystal structure of the MopR^AB^ in the phenol bound form shows that the binding pocket is buried inside the protein and there is no direct path to facilitate ligand entry (Figure 1b). Various attempts to crystallize the unliganded form of MopR^AB^ (Figure S1b) were unsuccessful, probably owing to the increased flexibility of the protein in absence of phenol. Therefore, we undertook MD simulations to achieve a conformation of the MopR sensor domain (MopR^AB^) (using PDB ID:5KBE) in its apo state.

Comparison of the equilibrated structure of apo with that of phenol bound states of MopR^AB^ reveals that although the phenol pocket still remains buried there are several other regions of the enzyme that show localized changes in structure. Maximum change was observed in two regions (Figure 1d): first, near the vicinity of the α2 (residues 48-62) and α5 (residue 138-147) and its following loop (residue 148-154) and the other being at the end of the sensor domain from α7 and loop preceding it (residues 194-212). The RMSD (Figure 1e) and RMSF plots (Figure S2) confirm that these changes are consistent through the simulation.

In the first region upon binding of phenol the helix α5 rotate away from α2 by about 20°, pulling the connector loops with it (Figure 1d). To gauge the significance of the associate motions this region was further subjected to a bioinformatic analysis which revealed that the NtrC subset of proteins that bind aromatic ligands (Figure S3, Table S1) contain a conserved glycine residue in the flexible ‘GAS’ motif (constituting of amino acids G148, A149, and S150 in MopR) (Figure S3, 1d) which resides at the tip of α5. Previous *in silico* studies have identified such small flexible residues like glycine, serine, alanine, as hinge regions and reports have attributed these motifs to be present where flexibility of motion is a prerequisite.^26–28^ The movements about strategic hinge regions have also been shown to play an important role in facilitating protein-ligand interactions or in domain movements to bring about allosteric communication.^27,29^ Using this premise, we referred to this region as the “G-hinge region” (Figure 1d,e). The second region where substantial motion was observed is the helix-loop motif that connects to the B-linker (Figure 1d,e). In the NtrC family of proteins, the B-linker serves as a connection point between the N-terminal sensor and the tandemly located ATPase domain.^14,15,20^ The sensing activity is attributed to be turned-on in the presence of the sensor molecule, phenol and the information is passed through this connector region.^15,21^ Comparison of MD snapshots of apo versus bound forms of MopR^AB^ (Figure S1b) show that the secondary structure in this region gets reorganized upon phenol binding (Figure 1d,e). Since this region is ∼20 Å away from the phenol binding site and a conformational change was nevertheless observed in response to phenol binding, the region has been referred henceforth as the “allosteric linker region” (Figure 1d).

To gain deeper insights into the process of how ligand binding propagates downstream signal relay, in-depth analysis of the apo and bound form of the structures was undertaken. It is a well-established fact that signal transduction in protein from one point to another propagates by gain and loss of a set of the interactions and several proteins operate via a large interconnected network of interactions.^30^ In order to unravel such an associated network of residues that bring about the communication of the allosteric linker and the G-hinge region, hydrogen bond propensities were calculated from the simulation trajectory (Figure 2,3,S4 Table S2-S4). MD results show that binding of phenol creates space in the binding pocket and drives a conformational change such that W134 and H106 shifts outwards (Table S5). For instance, H106 imidazole sidechain reorients to form hydrogen bonds with phenol and V112 (Figure 2a,b) and this motion in turn leads to decreased hydrogen bond propensities for the adjacent Q113(β4)-L135(β5) and M111(β4)-F138(α5) pairs that otherwise formed stable hydrogen bonds in the apo state (Figure 2a,b). This expansion of the binding pocket, upon phenol binding, percolates to α5 and results in its separation from α2 and α6 as represented by a rotation of ∼20° of α5 with respect to α2 in the apo verse the phenol bond forms. This is also exemplified by decrease in overall hydrogen bond propensities between residues on α5 with that of α2 and α6. For example, E139 of α5 with R55 of α2 and W156, M157, L158 of α6 (Figure 2c-f) all show mark shifts. The loss of E139-α6 and E139-α2 connections subsequently leads to disruption of hydrogen bond between H143(α5) and hinge residue G148 (Figure 2c,d). Therefore, pocket cap and the G-hinge region are linked through the above-mentioned residues, such that phenol induced conformational rewiring in the pocket cap that permeates to the G-hinge.

**Figure 2.**
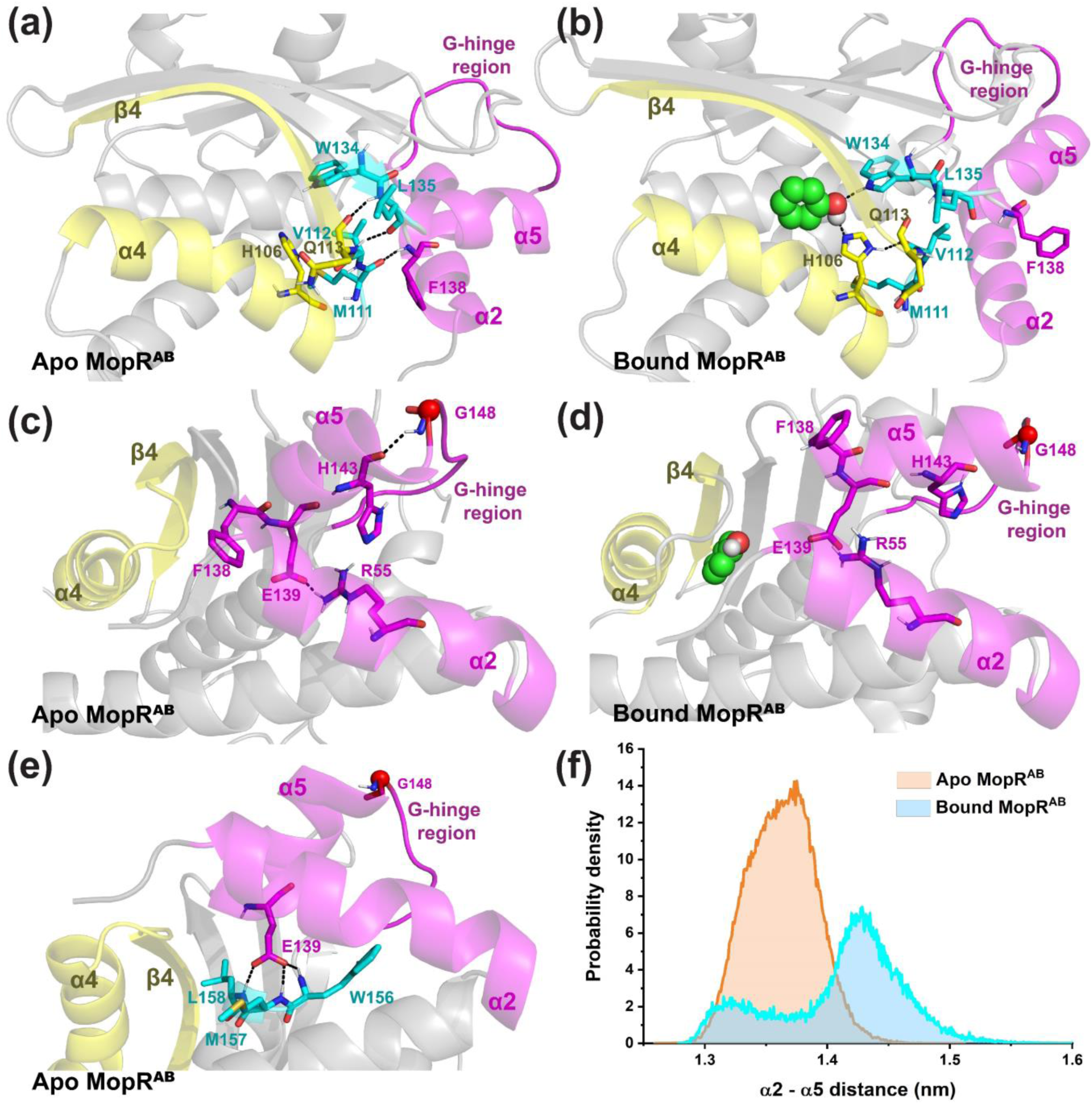
Signal transfer from phenol pocket to the G-hinge region. The panels **(a**,**c**,**e)** represent the predominant network that was observed during the MD trajectory based on hydrogen bond propensity analysis of apo MopR^AB^. **(b**,**d)** represents the hydrogen bond propensity analysis of bound MopR^AB^. **(f)** Probability density for the α2 and α5 distance over the last 500ns trajectory plotted.

**Figure 3.**
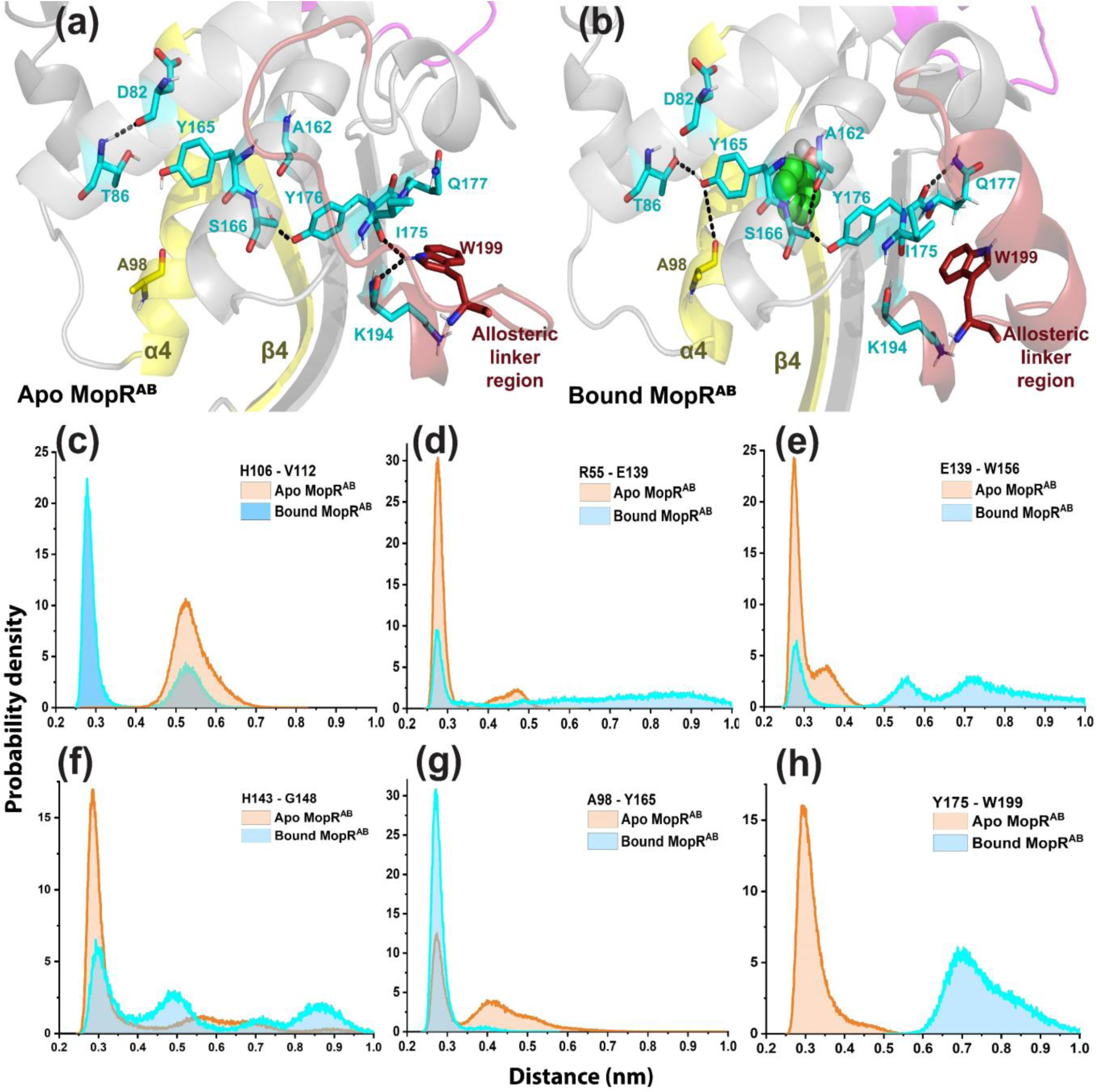
Hydrogen bond propensity profile for signal transfer from phenol pocket to the allosteric linker region. **(a)** and **(b)** represent the predominant network that was observed during the MD trajectory based on hydrogen bond propensity analysis of apo and bound MopR^AB^ respectively. **(c-h)** Probability density of inter-residue pairs for select residues whose hydrogen bond propensities significantly alter between apo and bound forms of MopR^AB^.

The mode of signal transfer from the phenol pocket to the allosteric linker region was also mapped. It was found that phenol binding also restructures this arm of the network. For instance, Y165 reorients to interacts with phenol, via aromatic stacking interactions, (Table S6) and this shift results in Y165 to form hydrogen bonds with A98 and T86 (Figure 3a,b). However, reorientation of Y165 affects other nearby interactions such as the hydrogen bond between D82 and T86 which was present in the apo state is now disrupted. Whereas, a new interaction network that connects the peripheral phenol binding pocket residue, S166, with both A162 and Y176 is formed. Y176 also gets reoriented via its backbone hydrogen bonding with amide group of Q177. It is noteworthy that Y176 acts as a bridging residue and upon phenol binding. Y176 additionally interacts with phenol through aromatic stacking interactions (Table S6).

The conformational rewiring in the above-mentioned binding pocket residues percolated to other adjacent residues, specifically a decrease in hydrogen bond propensities of W199 sidechain with I175 and K194 was observed (Figure 3a,b). The W199 consequently get released, hence relaxing the adjacent loop (202-210 of allosteric linker region) which then restructures into a helix as predicted by patterned change in the hydrogen bond propensities of C=O of i^th^ residue and NH of (i+4)^th^ residue (Figure 3a,b). Thus, the MD strongly suggests that sequential changes in the hydrogen bonding patterns allosterically connect the binding pocket and the allosteric linker region.

To summarize, the presence of phenol is percolated to the two extreme ends of the structure which is exemplified via changes in the hydrogen bond propensities. Apart from the predominant electrostatic effect that seems to govern the relay network in MopR, it was also noticed that an underlining subtle hydrophobic effect may play an integral role in effecting the relay. For instance, residue Y165 and Y176 that have strong shift in hydrogen bonding propensities upon phenol binding also form the wall of the phenol pocket and provide stabilizing stacking interactions that assist in phenol binding (Table S6). Hence, these residues support the relay network via dual contributions.

### Structural and thermodynamic probing of G-hinge region

To validate the MD results and discern the functional role of the G-hinge in ligand binding, we decided to mutate this region. It has been established that proline when introduced at the end of a helix has a destabilizing effect on the helix integrity.^31,32^ Moreover, in proteins that regulate the biosynthesis of aromatic compounds, the glycine residue has been shown to play an integral role in establishing a communication network between the active site and the distal amino acid binding regulatory site.^30^ This network was shown to be obliterated upon introduction of the G to P mutation in this system.^33^ Hence, with the two-fold objective of both restricting the local motion in this region and to change the capping residue on α5 from a stabilizing glycine to a destabilizing one, a G148P mutation was performed. The G148P mutant was expressed and purified and a comparison of the CD spectra of the G148P mutant (MopR^G148P^) with the wild-type protein (MopR^AB^) shows that the mutation does not cause perturbation in the overall secondary structure (Figure S5a,b). T_m_ studies also reveal that the thermal stability of MopR^G148P^ is comparable to that of the native protein (Figure S5c,d). Additional, size exclusion chromatography studies confirm that oligomeric state of the MopR^G148P^ is maintained as a dimer form in solution, similar to that observed for the wild-type (Figure S6), thus, this mutation does not perturb the structure in an adverse fashion. To discern if this mutation has any functional relevance, isothermal titration calorimetry (ITC) studies were carried out. Surprisingly, ITC studies show that MopR^G148P^ exhibits a 7-fold higher binding affinity compared to MopR^AB^ towards phenol (Figure 4a,b; Table S7) with a K_d_ value of (0.07 ± 0.02 μM). The data suggest that the effect of G148P substitution which is situated at a distance ∼20 Å percolates to the MopR^AB^ pocket, making the protein more conducive to phenol binding. The ITC corroborate our MD simulation results and reassert that there is indeed some communication between the phenol pocket and the G-hinge region. The G148P mutation likely results in a rearrangement of the network which is the dominant reason for the 7-fold increase in phenol affinity. Since the G-hinge dynamics seems to be correlated with the pocket we speculate that this region plays a role in either facilitating entry of the ligand or coordinated motions in this region that assist in phenol pocket formation.

**Figure 4.**
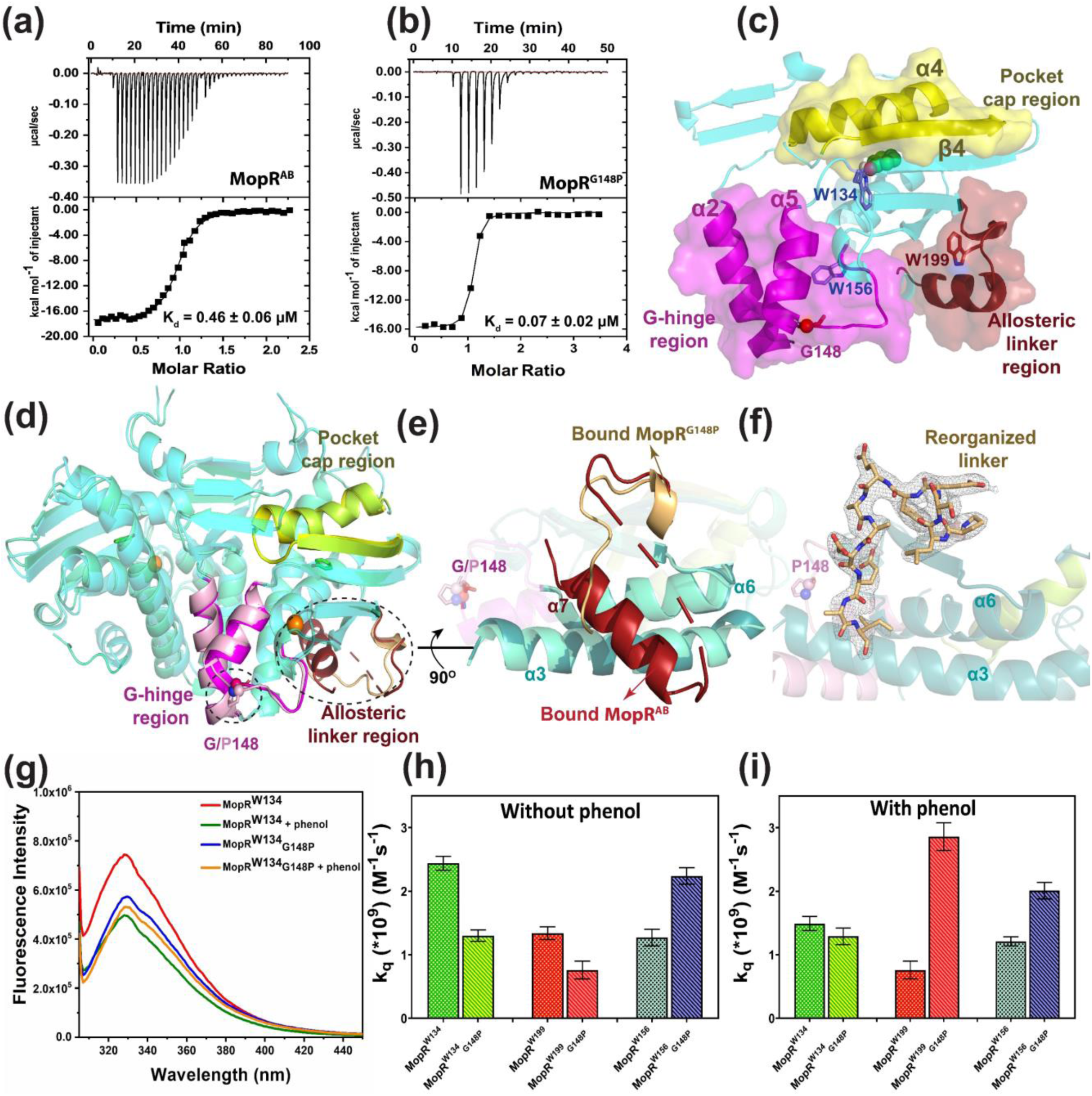
**(a**,**b)** ITC binding studies of MopR^AB^ (Ray et al., 2016) and MopR^G148P^ with phenol. The curves that correspond to raw data are shown in the top panel, and the curve fit in the bottom panel. Data were fit using one set of sites model and the thermodynamic parameters obtained from the curve fitting are given in Table S7. **(c)** Protomer highlighting three major allosteric regions and tryptophan residues used for fluorescence studies are highlighted as sticks. **(d)** Structural superposition of phenol bound MopR^AB^ (in cyan) and MopR^G148P^ (in aquagreen) highlighting the significant regions that show structural changes. **(e)** Magnified view depicting rearrangement in the B-linker region **(f)** shows the 2Fo-Fc map contoured at 1σ of the rearranged B-linker region of MopR^G148P^ (in stick representation). In all the panels, carbon atoms of phenol are depicted as green sticks and zinc as orange sphere. All the oxygen and nitrogen atoms are in red and blue respectively. **(g)** Steady state fluorescence spectra of MopR^W134^ and MopR^W134^_G148P_. Bar graph showing comparison of KI quenching constant (k_q_) for single tryptophan mutants, **(h)** in absence of phenol and **(i)** in the presence of phenol.

In order to further validate the role of the α5 region, we additionally attempted to crystallize the apo and phenol bound form of the hinge mutant MopR^G148P^, however, the crystals were only obtained in the presence of phenol. The structure of the sensor domain (Figure S1b) of MopR^G148P^in complex with phenol was determined to a resolution of 2.3 Å (PDB ID: 7VQF) by molecular replacement (MR) method using the native MopR^AB^-phenol structure as the search model (PDB ID: 5KBE). The data processing and refinement statistics are provided in Table 1. Analysis reveals that the overall structure of the sensor domain in the phenol bound form is similar with a RMSD of 0.63 Å aligning 180 residues (alpha carbon atoms) (Figure 4d). Comparative analysis shows that the introduction of a proline residue in G148P mutation indeed destabilizes α5 and induces its shortening such that F147 which was previously part of α5 now resides on a loop (Figure S7). Another notable observation from the crystal structure was a marked conformational change in α7 region that forms the allosteric hinge. Here, it was observed that residues 212-225, which fold as a helix (α7) in the native protein, is no longer visible. Rather this region becomes disordered and instead clear electron density for the region 201-207, which was previously disordered, can now be visualized (Figure 4d-f). The α7 is a part of the B-linker region that communicates the ligand binding to the AAA+ domain. Thus, it was surprising to observe that perturbation in the ‘GAS’ hinge which is separated spatially by more than 20 Å, causes a major rearrangement in the allosteric linker region. Since crystallography can only provide static snapshots, this result intrigued us to further probe protein dynamics in both apo and phenol bound forms.

**Table 1.** Crystallographic data statistics of MopR^G148P^ in the presence of phenol.

| Processing | MopR<sup>G148P</sup>+phenol<br>(PDB ID: 7VQF) |
| --- | --- |
| Wavelength(Å) | 1.54 |
| Resolution (Å) | 33.30-2.30 |
| Space group | C222<sub>1</sub> |
| Unit cell | a=62.20, b=118.30, c=56.70<br>$\alpha = \beta = \gamma = 90^\circ$ |
| Total reflections | 58593 (6957) |
| Unique reflections | 9592 (1131) |
| Multiplicity | 6.10 (6.15) |
| Completeness (%) | 99.6 (99.9) |
| I/$\sigma$(I) | 9.26 (2.29) |
| CC<sub>1/2</sub> | 0.99 (0.77) |
| $\dagger R_{\text{merge}}(\%)$ | 13.5 (88.2) |
| Refinement | MopR<sup>G148P</sup>+phenol |
| Total no. of non-hydrogen atoms | 1602 |
| Total no. of protein atoms | 1524 |
| Total no. of water atoms | 58 |
| Total no. of ligand atoms | 7 |
| Total no. of metal atoms | 1 |
| No. of reflections in refinement | 9591 |
| No. of reflections in test set | 480 |
| $R_{\text{factor}}(\%)$ | 23.2 |
| $R_{\text{free}}(\%)$ | 26.2 |
| Bonds (Å) | 0.002 |
| Angles (deg) | 0.414 |
| Most favored region (%) | 96.3 |
| Additional allowed region (%) | 3.74 |
| Disallowed region (%) | 0.00 |
| Mean B factors ($\text{\AA}^2$) for overall structure | 39.9 |
| $\dagger R_{\text{merge}} = \sum_{hkl} \sum_i \vert I_i(hkl) - \langle I(hkl) \rangle\vert / \sum_{hkl} \sum_i I_i(hkl)$. where $I_i(hkl)$ is the $i$th intensity measurement of reflection $hkl$, $\langle I(hkl) \rangle$ its average.<br>Values ( ) are for the highest resolution shell. | |

### Fluorescence studies to gauge protein dynamics

In order to understand the conformational heterogeneity in the apo versus the phenol bound forms of MopR^AB^ (Figure S1b) as well as changes induced on introduction of the G148P mutation, tryptophan fluorescence studies were undertaken. Three tryptophan residues naturally located respectively in the three strategic regions of the protein i.e., phenol pocket cap region, G-hinge region and allosteric linker region were used as markers to study the local environment for both the native and G148P mutant (Figure 4c).

For generating each single tryptophan construct, all other tryptophan residues were mutated to alanine or phenylalanine. The following constructs were made - MopR^W156^, MopR^W134^, MopR^W199^, MopR^W156^_G148P_, MopR^W134^_G148P_, MopR^W199^_G148P_ (Experimental Procedure and Table 2, superscript refers to the tryptophan residue retained). The structural integrity of all the mutants was tested and it was confirmed that it was maintained (CD spectra shown in Figure S8).

**Table 2:** Single tryptophan mutants of MopR^AB^ and their description.

| Mutations made | Final constructs |
| --- | --- |
| W37F_W199F_W156F | MopR<sup>W134</sup> (Active site) |
| W134A_W199F_W37F | MopR<sup>W156</sup> (Zinc site) |
| W37F_W156F_W134A | MopR<sup>W199</sup> (Linker site) |
| W37F_W199F_W156F_G148P | MopR<sup>W134</sup><sub>G148P</sub><br>(Active site with hinge) |
| W134A_W199F_W37F_G148P | MopR<sup>W156</sup><sub>G148P</sub><br>(Zinc site with hinge) |
| W37F_W156F_W134A_G148P | MopR<sup>W199</sup><sub>G148P</sub><br>(Linker site with hinge) |

Analysis of the steady-state fluorescence data, under saturating concentrations of phenol, reveals that for the native constructs, MopR^W134^ shows maximum fractional decrease in fluorescence intensity between the apo and phenol bound forms (Figure 4g, S9). This was not surprising as W134 is a key phenol pocket residue. This change in fluorescence intensity of MopR^W134^ could be attributed either to quenching observed as a result of phenol binding or due to restructuring of the phenol binding pocket. Steady-state studies were also performed with the hinge mutant MopR^G148P^ mutant that exhibits 7-fold enhanced phenol binding. Surprisingly, we observe that addition of phenol does not cause significant change in the fluorescence intensity of MopR^W134^_G148P_ (Figure 4g). Thus, if the quenching in MopR^W134^ was due to phenol binding, a similar effect should be observed in both the MopR^W134^ and MopR^W134^_G148P_, which was though not the case. Consequently, in MopR^W134^, the change in fluorescence can be attributed to a restructuring of the sensor pocket which is now primed for phenol binding and not an effect of direct quenching by phenol. Further, ITC derived thermodynamic parameters (Table S7) also show that the measured change in enthalpy (ΔH) upon phenol binding is lower in G148P mutant relative to the native MopR^AB^, reflecting lesser reorganization required for the pocket to accommodate phenol in the case of the mutant. In addition, the total entropy change associated with binding, ΔS, shows a net increase in the case of G148P as compared to native MopR^AB^, suggesting a preformed pocket for MopR^G148P^.

To reiterate, the fluorescence and the ITC data point to a more compact structure of the phenol pocket in the hinge mutant as compared to the native. To further corroborate the changes observed via our initial steady state studies, supporting fluorescence lifetime studies were also performed. The observations from the steady-state experiments (Figure 4g, S9), at all the three tryptophan positions in both native and G148P mutants, are largely reproduced in the mean fluorescence lifetime (τ_m_) (Figure S10 and Table S8). For instance, we observe a consistent decrease in τ_m_ upon binding to phenol in the MopR^W134^ variant whereas, this trend was not observed in the MopR^W134^_G148P_ mutant, reasserting that there has been an arrangement in the phenol pocket region upon introduction of the G148P mutation.

To obtain better insight into the conformational dynamics in the apo versus the phenol bound forms, lifetime studies were combined with KI quenching for both MopR^AB^ and MopR^G148P^ proteins (Figure 4h,i, S11). Due to the larger size as well as the charged nature of the iodide ion, it exhibits high polarizability and hence it cannot penetrate into the hydrophobic core. Thus, KI quenching studies provide a direct measure of the surface accessibility of a particular tryptophan residue via measurement of its k_q_. Table 3 and Figure 4 h,i list the values of k_q_ estimated for all the proteins with and without phenol. A significant decrease in the value of k_q_ was observed for MopR^W134^_G148P_ in comparison to MopR^W134^ construct indicating a decrease in solvent accessibility of the phenol-binding pocket upon introduction of the mutation. This reasserts that the apo form of MopR^G148P^ is more compact in the vicinity of the phenol binding region and this pocket has become less accessible to solvent. Evidence that the pocket is indeed preformed in the MopR^G148P^ is strengthened as k_q_ for both the apo and phenol bound forms were found to be similar in this mutant. Furthermore, the k_q_ was comparable to the phenol bound form of MopR^AB^ (Figure S1b) corroborating the fact that very little change in local environment occurs upon phenol binding.

**Table 3.** Bimolecular quenching constant (k_q_) values for the single tryptophan mutants estimated from mean lifetime (τ _m_) and Stern-Volmer constant (K_sv_).

| Protein | $\tau_m$ (ns) | $K_{sv}$ ($M^{-1}$) (KI) | $k_q$ ($\cdot 10^9$) ($M^{-1}s^{-1}$) (KI) |
| --- | --- | --- | --- |
| **MopR<sup>W199</sup>** | 3.24 ($\pm 0.07$) | 4.36 ($\pm 0.03$) | $1.34 \pm 0.06$ |
| **MopR<sup>W199</sup> + phenol** | 3.15 ($\pm 0.06$) | 2.39 ($\pm 0.08$) | $0.76 \pm 0.11$ |
| **MopR<sup>W156</sup>** | 3.59 ($\pm 0.05$) | 4.60 ($\pm 0.08$) | $1.27 \pm 0.06$ |
| **MopR<sup>W156</sup> + phenol** | 3.47 ($\pm 0.08$) | 4.21 ($\pm 0.06$) | $1.21 \pm 0.07$ |
| **MopR<sup>W134</sup>** | 2.70 ($\pm 0.02$) | 6.21 ($\pm 0.09$) | $2.30 \pm 0.05$ |
| **MopR<sup>W134</sup> + phenol** | 2.62 ($\pm 0.03$) | 3.90 ($\pm 0.08$) | $1.49 \pm 0.06$ |
| **MopR<sup>W199</sup><sub>G148P</sub>** | 2.68($\pm 0.02$) | 2.98 ($\pm 0.04$) | $1.11 \pm 0.04$ |
| **MopR<sup>W199</sup><sub>G148P</sub> + phenol** | 2.41 ($\pm 0.11$) | 6.90 ($\pm 0.11$) | $2.86 \pm 0.12$ |
| **MopR<sup>W156</sup><sub>G148P</sub>** | 2.03 ($\pm 0.05$) | 4.57 ($\pm 0.08$) | $2.24 \pm 0.08$ |
| **MopR<sup>W156</sup><sub>G148P</sub> + phenol** | 2.09 ($\pm 0.06$) | 4.21 ($\pm 0.07$) | $2.01 \pm 0.09$ |
| **MopR<sup>W134</sup><sub>G148P</sub>** | 2.79 ($\pm 0.03$) | 3.64 ($\pm 0.06$) | $1.30 \pm 0.05$ |
| **MopR<sup>W134</sup><sub>G148P</sub> + phenol** | 2.80 ($\pm 0.06$) | 3.62 ($\pm 0.07$) | $1.29 \pm 0.08$ |

The other tryptophan residues such as W156, which is in the vicinity of the α5 hinge region were also probed via KI quenching experiments. It was observed that W156 does not show any local environmental changes between the apo and phenol bound forms in the native protein. However, the G148P mutant has a different profile from that of the native where W156 is now more exposed to the solvent, as a result of the mutation. This we believe this a consequence of the helix α5 being destabilized in the G148P mutant, as observed in the MopR^G148P^ crystal structure. It is important to note that the changes in accessibility are observed only as the function of mutation and not due to phenol binding, suggesting that the mutation may have a global effect in altering the overall relay network.

The third tryptophan W199 resides in the allosteric hinge region. Snapshots from MD trajectory show that α7 region, on which W199 resides, shows a significant conformational change upon phenol binding. Moreover, since this helix adopts different conformations in both the phenol bound MopR^AB^ and MopR^G148P^ crystal structures, we envision this region to be dynamic. Since in other NtrC family proteins, B-linker is implicated in controlling ATPase activity via transmission of information between the signal sensing domain to the ATPase domain, via the B-linker, we believe that motions in the B-linker may have bearing on the de-repression mechanism. In MopR, we have exploited W199 as a reporter residue to gauge the conformational dynamics of this region. By observing the solvent accessibility of W199 in various states along with MD and structural data we aim to provide insights into the mechanism of communication.

Comparison of k_q_ values obtained from KI quenching studies show that W199 region is indeed dynamic and reorients upon phenol binding. However, the maximum difference in accessibility was observed between the phenol bound states of MopR^AB^ and MopR^G148P^. Comparison of the crystal structure of the MopR^AB^ versus MopR^G148P^ clearly shows that this region has a dramatic reorganization of the linker in both cases. While in MopR^AB^ the W199 is partially shielded by residues lining α7 in the G148P mutant this helix is disordered exposing the W199 to the solvent. This is supported by enhanced KI quenching as exhibited by a 3-fold increase in k_q_ in this mutant (Figure 5). Thus, fluorescence studies highlight that both the phenol pocket and the allosteric linker region are dynamic in nature and show different accessibilities both in the apo versus bound form. The changes in the G-hinge region are more subtle in nature and mutations in this region do not affect local environment, rather distal regions show differences if the G-hinge is perturbed.

**Figure 5.**
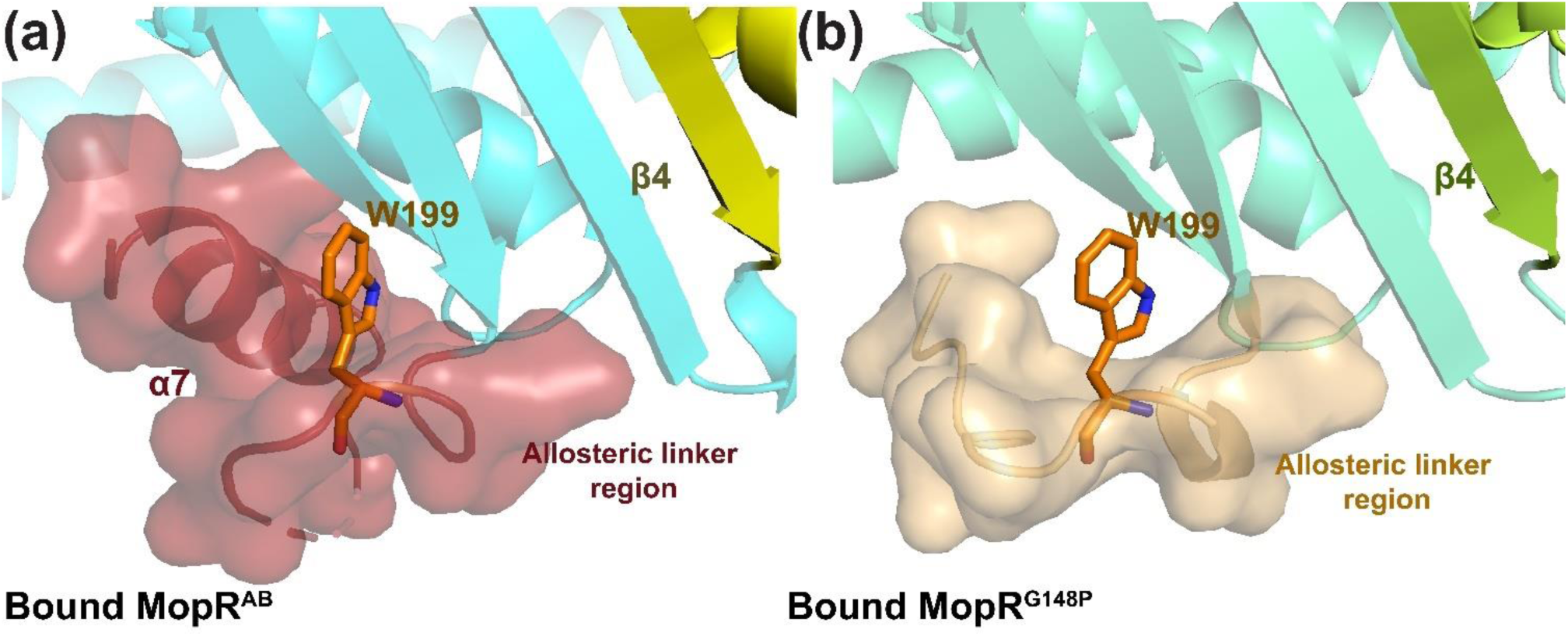
Flexibility of the allosteric linker. Environment of W199 (shown as orange sticks) in **(a)** bound MopR^AB^ and **(b)** bound MopR^G148P^. W199 is less accessible in native MopR^AB^ as compared to MopR^G148P^.

### Allosteric network and its implications to transcription activation

Mapping of the hydrogen bonding propensities obtained via MD simulations on the MopR^AB^ (Figure S1b) crystal structure helped create a communication network which connected the three strategic regions; phenol cap, G-hinge and allosteric linker. The electrostatic connections indicate that the G-hinge region likely communicates with the allosteric linker region via the phenol pocket cap (Figure 6a). The reliability of the hydrogen bond network was tested by creating mutations that disrupt the existing network. To access the role of this network in regulating the activity of the tandemly located ATPase motor, that is to understand the effect of these mutation on ATP hydrolysis, the mutations were introduced on the longer version of the MopR protein, MopR^A+C^ (Figure S1c), which comprised of both the phenol binding and ATP hydrolysis domain. All phenol affinities were measured only using the MopR^AB^ construct (Figure S1b). It was observed that any perturbation of the network results in marginal decrease in phenol affinity but leads to compete loss of the downstream ATPase activity. For instance, mutation of H106 and W134, where the G-hinge network originates, shows that although the H106A and W134A mutants are able to bind phenol with moderate affinity, a compete loss in ATPase activity occurs (Figure 6b). The observation highlights that it is not just the binding of phenol to the ligand pocket that determines downstream signaling, rather the contacts between the intermediatory residues are crucial for correct passage of the binding event. A similar scenario was observed for the phenol pocket-allosteric linker network, here also any perturbation emanating from the phenol pocket, completely abrogates ATPase activity as was observed for the proteins where the S166, Y165 and Y176 network has been individually disrupted via mutations of these residues (Figure 6b). It is noteworthy that in these mutants the phenol affinity is only marginally affected but ATPases activity shows a dramatic loss.

**Figure 6.**
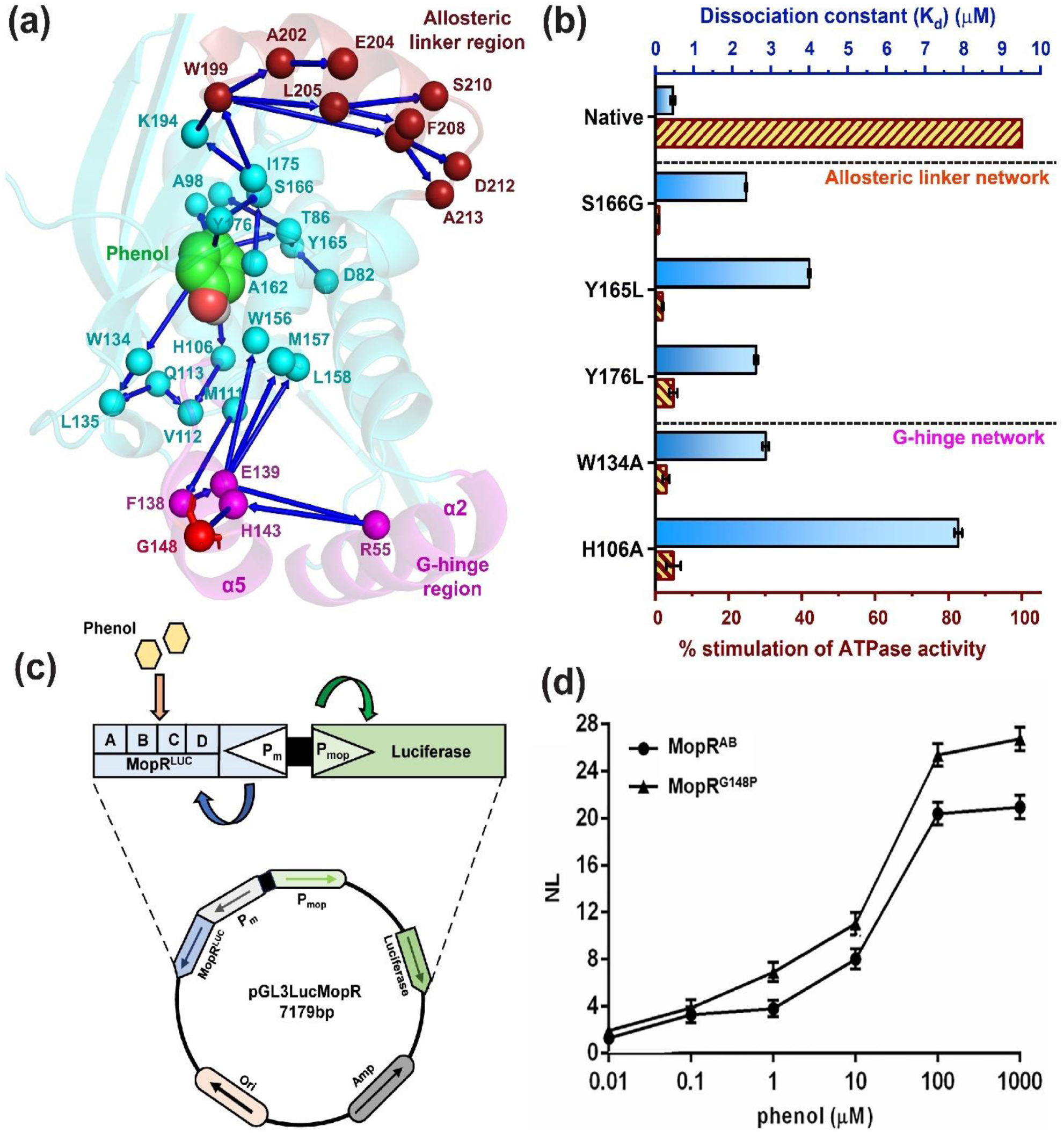
**(a)** Overall allosteric communication network in MopR sensory domain mapped using hydrogen bond propensities. **(b)** Bar graph representing the dissociation constant (K_d_) and % stimulation of ATPase activity of various mutants upon phenol binding. **(c)** Whole cell biosensor design of MopR based on a luciferase expression chemiluminescent readout. **(d)** Whole cell biosensing assay depicting enhanced luciferase activity for the MopR^G148P^ mutant (MopR^G148P^) as compared to the wild type.

The G148P mutation is at the end of the G-hinge network and here we observed that phenol binding had increased by 7-fold with no significant effect on the ATPase activity. Hence, to understand the effect and relevance of the mutation and its implications towards downstream transcriptional activity of MopR, an *in vivo* transcriptional assay was performed for both the native and G148P variant. A whole-cell setup (Figure 6c, S1d), where the entire upstream activation cassette, which includes the UAS, full-length MopR gene under its σ^70^ promoter and the downstream σ^54^ promoter was constituted and cloned in the pGL3basic vector (Figure 6c) such that the luciferase expression could serve as a reporter of the transcription activity (described in detail in experimental procedures). In *Acinetobacter calcoaceticus*, expression of MopR is essential for activation of the σ^54^-dependent RNA polymerase which then transcribes the phenol degradation cassette. Here, this gene cassette was replaced by luciferase to gauge the transcription efficiency of MopR and its G148P mutant. The transcriptional ability of both the native (MopR_LUC_) and mutated MopR (MopR^G148P^_LUC_) systems (Figure S1d) were subsequently tested using the entailed procedure. Briefly, the luciferase activity of both the constructs was recorded in the presence of varying concentrations of phenol. It was found that the luciferase signal of MopR^G148P^_LUC_ was always higher (∼25%) than the native MopR_LUC_ indicating better transcription ability of MopR^G148P^_LUC_ (Figure 6d). Thus, analyzing the results, it becomes apparent that the G-hinge mutant state is indeed effectively primed for downstream activation. It is not certain why the G-hinge mutant exhibits higher transcriptional activity or phenol binding. One reason we speculate is that G148P mutation facilitates the opening of the sensor domain and allows the phenol moiety to enter with more ease that subsequently expedites the downstream relay.

## DISCUSSIONS

Proteins elicit their function by shuffling between conformational states that allow for various important events such as binding of ligand to catalysis and in some instances facilitates transfer of information between active sites.^34^ In several of these systems, underlying allosteric regulation governs a switch between distinct states that allows for efficient function.^35^ Here, it is important to assert that allostery can operate both via inducing a major structural change, such as in conformational allostery, or in some instances subtle changes in side chain of certain amino acids, without any apparent structural change can contribute to function, like seen in systems that exhibit dynamic allostery.^9,36,37^ A common observation in both type of allosteric systems is a shift in hydrogen bonding network, where perturbation has resulted in dramatic change in the functional properties. For example, systems where conformational allostery is predominant, such as Hsp70, a complicated bi-directional hydrogen bonding network has been observed where allosteric cues between the nucleotide binding domain and the substrate binding domain are passed via rearrangement of this crucial connection.^38^ In this work, by employing MopR, as a model system, we explore the importance of allosteric networks that connect different segments of a protein domain and decipher how these connections, which sometimes are elusive and seemingly silent can cause changes in the protein network that have a profound effect on both affinity and activity. Two distinct networks, one that is primarily governed by dynamic allostery; the phenol pocket-G-hinge network and the other which exhibits aspects of both dynamic and conformational allostery namely the phenol pocket-allosteric linker network was unearthed.

The phenol pocket G-hinge network role was deciphered by examining the crystal structure of the phenol bound form and comparing it with the MD trajectory of the apo form. The X-ray structure shows that binding of phenol results in a compact structure where it binds to the interior of the protein, in a snug pocket. Further, MD simulations of the apo form shows that the structure in the absence of phenol is more or less similar and only few notable conformational changes were observed. However, a noteworthy feature was a distinct shift in the electrostatic network that respectively connects the G-hinge and the allosteric linker regions with the phenol pocket. For instance, it was observed that phenol binding induces a shift in the hydrogen bonding propensities of W134 and H106, this effect percolates to the next residue in the relay and finally the network that terminates at the G-hinge which is different from the apo state. What was most fascinating is that an analysis of the residues that partake in the relay network reveals that the residues are more or less conserved in close homologs of MopR such as PoxR, DmpR, XylR, etc (Figure 7). It was observed that the side chain residues that form an integral part of the network were completely conserved whereas, residues which contribute through their backbone show some variation. This highlights the fact that the network is more like a community network which has co-evolved in these members such as to allow a more orchestrated shift between distinct allosteric states that are functionally relevant. Perturbation of this network by introducing a G148P mutation experimentally confirmed that indeed the phenol pocket is influenced by changes in the G-hinge region. Here, we think that the G-hinge network most likely assists in entry of phenol as MD simulations hint that, synchronous fluctuations between the G-hinge region and the pocket cap may facilitate phenol binding. In the apo state the protein likely accesses multiple conformational states and it is possible that the state conducive for phenol entry, where coordinated motion between the pocket cap and the G-hinge occurs, allows for effective phenol access. The observation that the G148P mutation affects phenol affinity and corroborating fluorescence studies which show that the phenol pocket is partially pre-formed in G148P, confirms that perturbation in the G-hinge is noticed at the phenol binding site. These observations reassert that G-hinge and phenol pocket region are strongly connected. However, introduction of G148P does not cause any perturbation in the transmission network, as relay is not disrupted. Supporting transcription activity studies for this mutant further corroborate that this mutant not only aids in phenol binding but also exhibits an enhanced overall transcriptional rate of around 25%. Thus, it appears that by introducing the G148P mutation, we have shifted the conformational landscape of MopR and it predominantly now accesses states that are more conducive to allow for quick passage of signal that favor downstream function.

**Figure 7.**
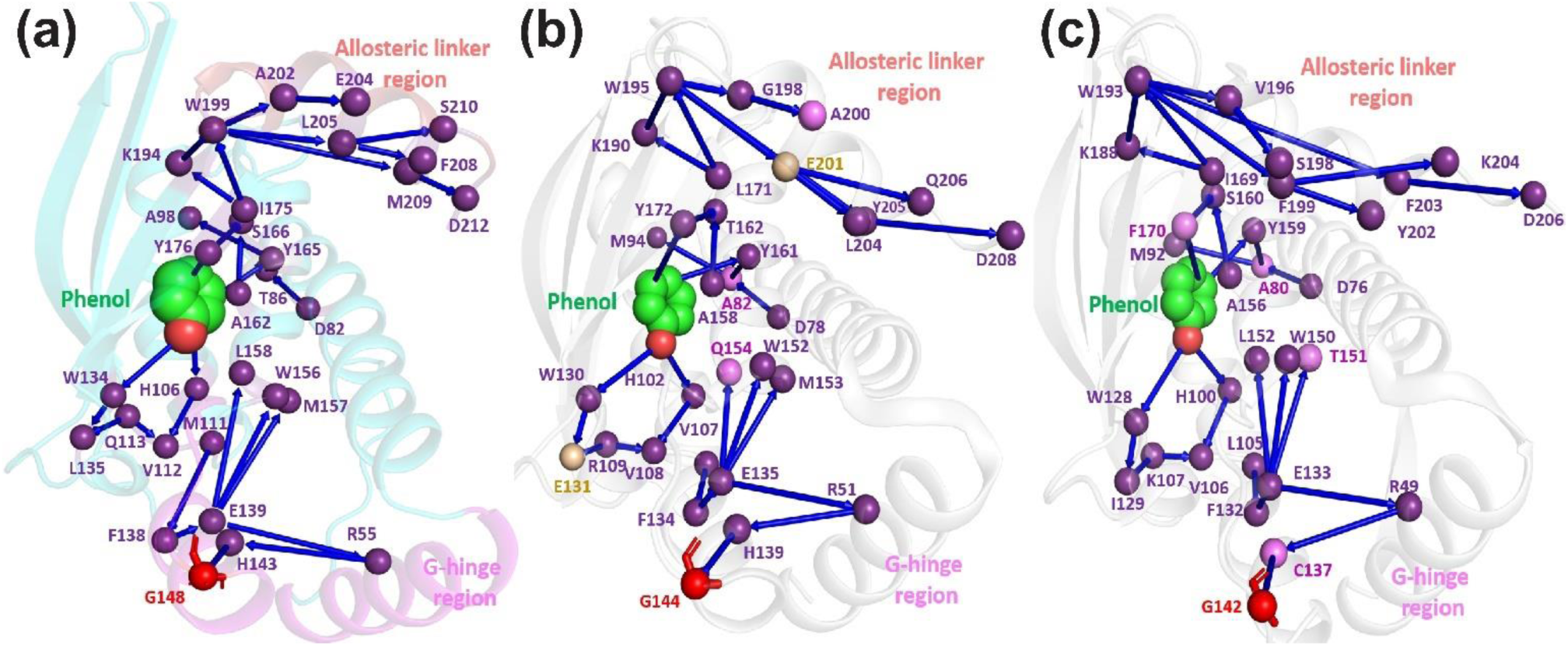
Representation of community networks in aromatic compound sensing NtrCs. The networks were constructed based on similarity of sequence and hydrogen bond connections visualized from available crystal structures of **(a)** MopR (PDB ID: 5KBE) **(b)** PoxR (PDB ID: 5FRW) **(c)** DmpR (PDB ID: 6IY8). Violet sphere, pink and yellow represent identical/similar, slightly non-identical and completely non-identical amino acid residues. Hinge glycine residue is shown in red.

The network established for MopR protein is primarily governed by rearrangement of the electrostatic interactions and only subtle structural changes are observed therefore, it appears that dynamic allostery is predominantly at play. Comparative analysis of thermodynamic parameters of MopR in both the native and G148P mutant shows that both the enthalpic and entropic contributions influence binding. It was earlier envisioned that in systems that exhibit dynamic allostery such as, catabolite-activating protein (CAP) and PDZ domains entropy-driven allostery solely governs conformational fluctuation.^7,39^ However, recent reports by Kumawat et al. clearly show that dynamic allostery can encompass enthalpic effects.^9,40^ In their work, they show that restructuring of the electrostatic network in the PDZ protein has an enthalpic contribution. Deletion of a helix that lies at the periphery of the PDZ domain does not perturb the overall structure but it results in 21-fold reduction in affinity of the ligand.^39^ This is because this helix assists in a rearrangement of the electrostatic network upon ligand binding. On a similar vein, in MopR, introducing a G148P mutation likely alters the overall phenol binding dynamics of MopR^AB^ resulting in a 7-fold increase in affinity for this mutant. Here, also both entropic and enthalpic contributions were found to be key players that determine the shift. Thus, dynamic allostery in the MopR system governs the communication between strategically located hotspots in this protein. Such type of changes in affinity via the dynamic allosteric pathway are also observed in cold adaptation of enzymes, as observed in adenylate kinase, where specific distal surface mutation that increase the propensity of the enzyme to unfold, result in marked change in ligand affinity.^41^

Apart from the phenol pocket-G-hinge network, the other network that also originates at the phenol pocket and terminates near the end of the ligand binding domain, near the B-linker region also showed interesting features. This phenol pocket-allosteric hinge communication network is an example of a complex interplay of conformational and dynamic allostery. The B-linker helix at the end of the phenol binding domain connects this domain with the tandemly located ATPase domain and it is envisioned that entry of phenol stimulates ATPase activity of the AAA+ domain via conformational changes in the B-linker that are passed on.^20,21^ Therefore, dynamic nature of the B-linker as corroborated by fluorescence, and X-ray crystallographic studies was expected. What is more interesting is the presence of dynamic allostery that is facilitated via a conserved network of connections that run between the phenol pocket and tip of the B-linker that help communicate the progress of phenol binding. Comparison of the apo and bound forms of ligand binding domain show that a rewiring of the electrostatic connections between the phenol pocket and the B-linker in the presence of phenol enables passage of information. Thus, the ATPase domain is informed of phenol binding via dynamic allostery within the internal network that spans within the phenol binding domain and expressed as conformational allostery via structural motion in the B-linker. Both, ITC binding studies and ATP hydrolysis studies with variants of MopR, where this network has been disrupted show the importance of conserved interactions for effective relay. Almost complete loss of ATP stimulation is observed whenever the network is disrupted. In some of these cases, such as mutations at the periphery of the phenol pocket, as in W134A and H106A, we observe that these point mutations are still capable of binding phenol, although with somewhat decreased affinity. However, the ATP stimulation is completely obliterated due to these mutations. Other mutations in the relay that are not directly involved in phenol binding leave the phenol binding almost unaltered but again complete loss in ATPase activity occurs. This highlights the importance and precision with which the relay network has to operate to elicit function and asserts the idea that phenol binding is not the rate determining step that controls MopR function, rather it is the integrity of the relay that is paramount to maintain function. The importance of this network is again highlighted as it was observed that like in the case of the G-hinge network, the phenol pocket-allosteric hinge relay network is also conserved in other proteins such as PoxR, XylR and DmpR that bind similar aromatic compounds.

The MD simulations indicated independent relay networks emanating from the phenol pocket to the G-hinge and allosteric linker. However, a striking observation that emerged from the crystal structure of the ligand binding domain of G148P mutant was a rearrangement of the B-linker. This was unexpected as MD analysis does not reveal any direct connection of the G-hinge with the allosteric linker region. Thus, this experimental observation seeded the idea that the phenol pocket, G-hinge and allosteric linker are all connected and the networks operate in an interdependent fashion. It can be envisioned that the networks run via the central phenol pocket and are perceived by other functionally important parts of the protein. ITC and transcriptional activity assay also confirm this idea, otherwise a mutation in the G-hinge would not have such far reaching functional effects. The study emphasize that it is not only the local active site region that partakes in function rather seemingly distal and benign parts of protein domains have a profound influence on functional and binding properties of proteins. It reasserts that apart from the obvious active site residues several other portions of the protein also play an integral role in determining the functional outcome of an enzyme.

In conclusion, here we show via a combination of MD and experimental approaches that MopR has distinct regions that form a highly involved bi-directional network. The central phenol pocket within the ligand binding domain connects to two other regions G-hinge, that likely influence entry of phenol, and allosteric hinge, that passes the signal to the downstream ATPase domain. Phenol binding induces a rewiring of the electrostatic connections that brings about function by eliciting dynamic allostery. It was established that it is not the ligand binding that determines downstream ATP hydrolysis, rather the order in which the rearrangement of interactions that occur upon ligand binding is more crucial in maintaining the integrity of the relay. Further the flexibility of the G-hinge was probed by undertaking a G148P mutation and it was determined that G-hinge influences both phenol binding as well as downstream transcription. Changes in this region are communicated via the bi-directional network that spans across the ligand binding domain. The work highlights the importance of long-distance communication and shows how relay networks in proteins are highly sensitive to disruption and an important backbone that support protein function.

## MATERIALS AND METHODS

### Molecular Dynamics Simulations

The computational structural model for the phenol bound wild type (Bound MopR^AB^) was the crystal structure of a dimer of MopR sensor domain (PDB ID: 5KBE) (Fig. 1B, S1B). For apo simulations, the coordinates of phenol ligand were removed from this crystal structure (Apo MopR^AB^). All the crystal water molecules were removed. The missing crystal residues were modelled using CHARMM-GUI.^42^ The N- and C-terminus of the protein were capped by NH^3+^ and COO^-^. The protonation state of all amino acid residues corresponds to neutral pH except Zinc coordinated cysteines (C155, C181, C189) which were modelled in their deprotonated states.^43^ The proteins, the ligand (phenol) and the ions were modelled using Charmm36m forcefield.^44^ The systems were solvated with using the TIP3 water model^45^ where for each of the system state, the starting models were placed at centre of truncated octahedron box with empty box volume filled by TIP3P-charmm water (∼18500 molecules) and neutralized by 200 mM NaCl ions. The system was energy minimized using steepest descent algorithm with position restraint on all protein heavy atoms. After minimization, 500 ps position restrained MD simulation is performed in NVT ensemble at 298 K temperature maintained by Nose-Hoover thermostat^46,47^ with relaxation time of 1 ps. The system was subsequently subjected to unrestrained MD simulation for a duration of 1µs in NPT ensemble at 298 K temperature (using Nose Hoover thermostat) and 1 bar pressure using Parrinello-Rahman barostat^48^ with a coupling constant of 5 ps. All simulations were performed with 2 fs timestep using Leap-frog integrator with-in GROMACS 20XX simulation package. Periodic boundary condition was implemented in all three dimensions. The Verlet cut-off scheme^49^ was employed for Lennard Jones interaction and short-range electrostatic interactions. Long range electrostatic interactions were treated by Particle Mesh Ewald summation method.^50^ All hydrogen bonds were constrained using LINCS algorithm.^51^ The bonds and angles of TIP3P water molecules were constrained using SETTLE algorithm.^52^ The simulations with apo protein were replicated in three independent trajectories. The final 500 ns of each of the 1 µs long trajectories were considered for atomistic analysis.

### Molecular Dynamics Simulations analysis

For each of the system state, hydrogen bond profiles were analysed using a criterion of distance ≤ 3.5Å and angle ≤ 30°.^53,54^ The propensity of hydrogen bond formation is calculated for individual hydrogen bonds as time average over all the sampling for each system. The propensity of hydrogen bond formation is basically the probability of finding a hydrogen bond between two residues and ranges between 0 and 1. The hydrogen bond propensities showing difference ≥ 0.5, were considered significant. The significant difference is considered if it follows either or all of the criteria. The first criteria considered was if hydrogen bond propensity showing difference ≥ 0.5. The second criteria was if a residue has decreased hydrogen bond propensity with one residue and simultaneous increase in hydrogen bond propensity with other residue as system go between different system states, with total difference ≥ 0.5.The third criteria being a residue has decreases/increased hydrogen bond propensity with multiple residues with total difference ≥ 0.5.

π-π aromatic stacking was considered for residues tyrosine, tryptophan, phenylalanine and histidine^55^ if the stacking pairs, are within 7 Å distance^54^. In order to calculate the pocket volumes, MD pocket utility is used.^56^ The pairwise distance between α2 (residues 48-62) and α5 (residues 138-147) was determined by taking the average distance of all possible pairs of mainchain atoms.

### Bioinformatic analysis

For mapping the sequence conservation, sequences of NtrC family proteins that are specific to the aromatic hydrocarbons were chosen. Details of the proteins are given in Table S1. The sequence alignment was performed using ClustalW.^57^ The sequence logo was generated using WebLogo.^58^ The sequence conservation was estimated as bits score on Y-axis and residue number on X-axis. A value of ∼4.3 corresponds to strict identity of the amino acid residue.^58^

### Site-directed mutagenesis and protein purification

The recombinant pET vector construct of MopR^AB^ (Fig. S1B) was used as template^14^ to make the glycine to proline (G148P) hinge mutant, MopR^G148P^. For fluorescence spectroscopy studies, the following mutations were made to generate single tryptophan constructs using MopR^AB^ (Fig. S1B) as the starting template: W134A_W199F_W37F (MopR^W156^), W37F_W156F_W134A (MopR^W199^), W37F_W199F_W156F (MopR^W134^), W134A_W199F_W37F_G148P (MopR^W156^_G148P_), W37F_W156F_W134A_G148P (MopR^W199^_G148P_), W37F_W199F_ _G148P_ W156F_G148P (MopR^W134^_G148P_) (Table 2). For mutants Y165L, Y176L, S166G mutants, the sensor domain constructs were generated using MopR^AB^ (Fig. S1B) as template and construct MopR^A+C^ (Fig. S1C) constituting of both sensor and ATPase domain using a template that encodes 1 to 500 residues of the *mopR* gene.

All the mutants were made by employing standard site-directed mutagenesis protocol using the Phusion DNA polymerase from New England Biolabs.. The mutant expression constructs were subsequently transformed into *Escherichia coli* BL21(DE3) plysS cells, over expressed with 1 mM IPTG (isopropyl-β-D-thiogalactopyranoside) as six-his tag fusion proteins and cultured at 16ºC for 16 hrs. All the mutated proteins were purified using Ni-NTA resin by standard His-tagged affinity purification protocol. The composition of the buffers used in subsequent purification steps was as follows: lysis buffer (50 mM Tris-HCl buffer, pH 7.5; 2 mM imidazole; 200 mM NaCl), wash buffer (50 mM Tris-HCl buffer, pH 7.5; 30mM imidazole; 200 mM NaCl), and elution buffer (50 mM Tris-HCl buffer, pH 7.5; 350 mM imidazole; 100 mM NaCl). The eluted fractions were desalted using an Econo-Pac 10DG (Bio-Rad, CA, USA) column that was pre-equilibrated with a desalting buffer containing 25mM Tris-HCl buffer, pH 7.5; 80 mM NaCl, 5% glycerol, and 0.5 mM DTT. The desalted protein fractions were pooled and concentrated up to 5-8 mg/ml. The fractions were then flash-frozen in liquid N_2_ and stored at −80 °C until they were used. The purity of the protein was verified by running a 10% SDS-PAGE followed by Coomassie Blue (HiMedia, Mumbai, India) staining.

### Ligand-binding experiments using ITC

All the ITC experiments were carried out on the constructs of MopR^AB^ constituting of residues 1 to 229 (Fig. S1B). All the protein and ligand samples were prepared in a buffer that contained 25 mM HEPES (pH-7.5) and 80 mM NaCl. In the ITC experiment, phenol was titrated against buffer and subtracted from the raw data prior to model fitting, in order to nullify the heat of dilution. 40 μM of MopR^AB^ and 10 μM of MopR^G148P^ was titrated with 400 µM and 200 µM of phenol respectively. For the Y165L, S166G and Y176L mutants, 25 μM of protein and 400 μM of phenol were used. The K_d_ values for W134A and H106A were taken from the previously reported results.^14^ The volume of the titrant (ligand) added at each injection into the sample cell was 2 µl for 5sec. A set of 20 injections were performed for each experiment with an interval of 120 sec between each successive injection. The temperature was maintained at 25 °C. The stirring rate was kept constant at 1000 rpm throughout all the ITC experiments. The data obtained were fitted and analyzed with Origin 7 software using ‘one set of site model’^59^. The curve fitting was done in the acceptable experimental window of c values of 10 ≤ c ≤ 500, where, c-value = n[Protein]/ K_d_ for n non-interacting identical sites (K_d_ is the dissociation constant).^60^

### Size Exclusion Chromatography (SEC)

A Superdex 75 10/300 GL column installed on an NGC Chromatographic System (BioRad) and equilibrated with the sample buffer (25 mM Tris–HCl (pH 7.5), 80 mM NaCl, 5% glycerol and 0.5mM DTT). 100 µL of ∼4mg/mL MopR^AB^ and ∼2mg/mL of MopR^G148P^ in sample buffer was injected onto equilibrated column at 4 °C at a flow rate of 0.3 ml/min. The column was calibrated with the following standards, carbonic anhydrase (29 kDa), Ovalbumin (44kDa) and Conalbumin (75kDa).

### Crystallization of MopR^G148P^

The purified his-tagged MopR^G148P^ (10 mg/ml) was first screened for crystallization using several commercially available crystallization screens such as Crystal screen, PEG/Ion (Hampton Research) and JCSG suite, PACT suite (Qiagen) using a Crystal Pheonix crystallization robot and the sitting-drop vapor diffusion technique at the crystallization facility at IIT Bombay; however, no notable crystal hits were found in any of the screens. Attempts were then made to co-crystallize MopR^G148P^ with phenol. MopR^G148P^ was incubated at 4 °C for around 30 minutes with 5 mM of phenol and was then subjected to crystallization trials in a manner like that of apo protein, using the screens that are commercially available at Hampton and Qiagen. All the protein-ligand complex solutions were filtered before crystallization trials. Crystals were obtained within seven days of setup in the following condition 0.2 M Magnesium acetate.4H_2_O, 15% w/v PEG 3350. The crystals were further optimized at 20 °C using the hanging-drop vapor-diffusion method, with 1μl of a protein solution, 1 μl of a precipitant solution, and a 500 μL reservoir volume. The trays were monitored in a temperature-controlled cabinet. Diffraction-quality isolated crystals of the maximum size grew (100×70×70 μm3) over a period of 10-12 days. Under optimized conditions, MopR^G148P^ in complex with phenol crystallized in the C-centred orthorhombic space group C222_1_ with unit cell dimensions of a=62.20, b=118.30, c=56.70 and α=β=γ=90°.

### Data collection and processing

X-ray diffraction experiments were performed at the home source of Indian Institute of Technology (IIT) Bombay using a Rigaku MicroMax-007HF X-ray diffractometer. A single crystal of the ligand complex was cryo-protected with 20%(v/v) ethylene glycol (prepared using mother liquor) prior to data collection. The crystal was then flash cooled in liquid nitrogen and transferred to a stream of nitrogen gas at 100K. X-ray data were collected at a wavelength of 1.5418Å on a Rigaku R-AXIS IV++ detector. The dataset was indexed, integrated and scaled with XDS.^61^ Data-collection statistics are summarized in Table 1.

### X-ray crystal structure solution

The structure of MopR^G148P^ co-complexed with phenol was determined at 2.3 Å by the molecular replacement method using the molecular replacement module of the PHASER program^62^ and native monomeric unit of MopR^AB^-phenol complex structure as a search model (PDB ID: 5KBE). Manual model building of the partially refined structures was carried out using the graphics program COOT^63^ and they were further refined using REFMAC5^64^. All Figures were made in PyMOL.^65^

### Circular Dichroism (CD) studies

CD spectra of the MopR^AB^, MopR^G148P^ and single tryptophan mutants were recorded on a Jasco J-815 CD spectrometer. The protein concentration used were 0.5 mg/mL and phenol used was 0.2 mM respectively. All the protein and ligand samples were prepared in phosphate buffer (25 mM sodium phosphate pH 7.5, 80 mM NaCl). Using same method, the CD spectra was recorded for all single tryptophan mutants (Data shown in Fig. S8). Scans were performed at 20°C using 0.1 cm path length quartz cuvettes with 8 sec differential integration time at a scan rate of 50 nm/sec. The mean residual ellipticity (MRE) in units of deg.cm^2^.dmol^-1^ was determined using the formula^66^,

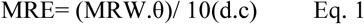

where θ is the observed ellipticity (degrees), d is the pathlength (cm) and c is the concentration (in units of g/ml). The MRW is obtained by dividing the molecular mass by N - 1, where N is the number of amino acids. The CD based thermal denaturation experiment were performed for MopR^AB^ and MopR^G148P^ to determine the stability of these proteins. Scans were performed at a temperature range of 20°C to 95°C using 0.1 cm path length quartz cuvettes with 16 sec differential integration time at a scan rate of 100 nm/sec with 3 mins delay time per temperature change.

### Steady state fluorescence measurements

Absolute fluorescence intensities were measured on FluoroMax-4 spectrofluorometer with 4 μM of protein samples in buffer (25 mM HEPES, 80 mM NaCl). The quenching experiments were performed on a Varian Cary Eclipse spectrofluorometer with 12 μM of protein in buffer (25 mM HEPES, 80 mM NaCl) titrated with increasing concentration (0-300mM) of potassium iodide (KI). For phenol bound form, the mutants were incubated with 5 mM phenol for 30 minutes prior to study. All the experiments were performed in quartz cuvette of 1 cm path length and the samples were excited at 295 nm and the emission spectra were recorded for 300 to 450 nm wavelength range. All measurements were carried out in triplicates and mean ± SE (Standard error) have been reported.

### Time-resolved fluorescence measurements

The time-resolved fluorescence decay was recorded using Ti-sapphire laser (Mai Tai HP, Spectra Physics) pumped by an Nd: YVO4 laser (Millennia X, Spectra Physics) generating the 885 nm pulses of width ∼1 ps. A flexible second- and third-harmonic generator (GWU, Spectra Physics) was used to obtain the frequency-tripled laser of 295 m for excitation. Fluorescence emission was collected through a 305 nm cut-off filter to exclude scattered photons completely when the monochromator was set at 335 nm. To obtain fluorescence lifetimes, a polarizer oriented at the magic angle (54.7°) was used to eliminate anisotropy decay artifact in the fluorescence decay data. All the measurements were made in 1 cm path length with 10 μM concentration of protein. All measurements were carried out in triplicates and mean ± SE (Standard error) have been reported.

### Fluorescence data analysis

The obtained decay curves collected at the magic angle were deconvoluted with the IRF (Instrument response factor) by using nonlinear least-square iterative deconvolution method based on the Levenberg–Marquardt algorithm and expressed as a sum of exponentials with equation,

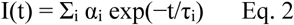

where, I(t) is the fluorescence intensity at time t with αi being the amplitude of the i^th^ lifetime τ_i_ such that Ʃ_i_ α_i_ = 1. The average fluorescence yield was estimated by calculating the mean lifetime using the equation: τ_m_ = Ʃ_i_ α_i_ τ_i_. The goodness of fits was assessed from the reduced chi square (χ^2^) values as well as from randomness of the residuals.

For monitoring the solvent accessibility of the four tryptophans, the KI quenching experiments were done and fitted to Stern-Volmer equation^67^:

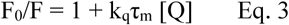

Where, F_0_ and F are the fluorescence intensity in the absence and presence of quencher KI respectively, [Q] is the concentration of quencher and k_q_ is the bimolecular quenching rate constant (M^−1^ s^−1^). τ_m_ is the mean lifetime in the absence of Q. K_sv_ is the Stern Volmer constant given by k_q_τ_m_ and is obtained from slope of Stern Volmer plots.

### Colorimetric ATPase assay

The Malachite green assay was used for determining the ATPase activities. The assay was carried out on the constructs of MopR^A+C^ constituting of residues 1 to 500 (Fig. S1c). 2µM of protein sample was incubated with 1µM phenol. 2mM of ATP was then added to the reaction mixture, and the reaction was allowed to proceed for 15 minutes followed by quenching with 0.5M EDTA. The dye reagent (consisting of Malachite green, Ammonium heptamobolybdate and Tween20) was added to the reaction mixture to form a phosphomolybdate complex zwhich gave a green color and for which absorbance was measured at 630 nm. All the data was collected in triplicates to estimate the errors.

### Construction of the whole cell MopR_LUC_ and MopR^G148P^_LUC_ construct

The whole cell construct of native MopR was first designed (MopR_LUC_) which consists of the full length *mopR* gene under the control of *P*_*m*_*-P*_*mop*_ promoter with a *luc* reporter module attached upstream, cloned into a pGL3basic expression vector that has been purchased from Promega (WI, USA) (Fig. S1D). *P*_*m*_ is the σ70 based promoter that controls the *mopR* gene transcription activity and *P*_*mop*_ is the σ54-based promoter which triggers downstream catabolic pathways on activation of MopR with suitable pollutants. For cloning experiment, firstly, the purified genomic DNA of *A. calcoaceticus* NCIB8250 (2μg/μL)^16^ was used as a template for the PCR amplification of *mopR - P*_*m*_*/P*_*mop*_ gene sequence. PCR was performed by using forward primer 5’-ACCGAGGTACCATTTAAGCCCGA-TAATTTA A-3’ and a reverse primer, 5’-ATTGACTCGAGATTCCGCTC ACCAGTAATAC-3’, with XhoI and KpnI sites at the ends. The amplified gene was then cloned into the pGL3basic vector using standard cloning techniques (Fig. 6c). This native whole cell clone was further used as a template to make the G148P mutant (MopR^G148P^_LUC_) in *mopR - P*_*m*_*/P*_*mop*_ by employing standard ‘site-directed mutagenesis’ protocol using the “site-directed mutagenesis kit” from Kapa biosystems. The cloned MopR^G148P^_LUC_ construct was then transformed into *Escherichia coli* DH5α cells and pre-inoculum was setup at 37°C overnight with the transformed colonies for performing the luciferase assay.

### Luciferase assay design on MopR_LUC_ and MopR^G148P^_LUC_

The luciferase assay described in this work is based on the luciferase assay system kit protocol from Promega (WI, USA) which was performed to monitor the downstream transcription activation potential of the native and mutated MopR.^68^ The overnight grown cultures of *E*.*coli* DH5α cells harboring the recombinant MopR_LUC_ and MopR^G148P^_LUC_ plasmids were sub-cultured and grown to the log phase till the OD at 600 nm reached ∼0.3. This was followed by supplementation with phenol at a gradient concentration of 0.1-1000 µM. After 1 hour of induction, 50 µl of the cells were withdrawn, and frozen at -70°C with the addition of 5 μl of 1 M KH_2_PO_4_ and 20 mM EDTA (pH 7.8). The cells were lysed by the addition of 150 μl of lysis solution (1.25 mg/ml lysozyme, 2.5 mg/ml BSA, 1X CCLR) (Promega, WI, USA) and incubating at room temperature for 10 min. Supernatants were obtained by centrifugation. For the luciferase activity, 20 μl of supernatant was mixed with 50 µl of firefly luciferin solution (Promega, WI, USA). The bioluminescence was measured for 30 sec by Luminometer Berthold Detection System (Germany). Induction of the MopR_LUC_ and MopR^G148P^_LUC_ by phenol was expressed as normalized luminescence (NL) calculated as follows:

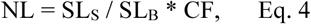

where SL_S_ is the luminescence of the biosensor in the dilution of the aromatic pollutant, SL_B_ is the background luminescence of the sensor bacteria, and CF is the correction factor.

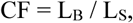

where L_B_ is the background luminescence of the control bacteria, and L_S_ is the luminescence of the control bacteria in the presence of an aromatic compound. The detection limit for the sensor bacteria for different pollutants was defined as a concentration of the compound which induced the sensor twice above the background level, i.e. NL ≥ 2. All data were collected in triplicates.

## ACKNOWLEDGMENTS

We wish to thank Prof. Jayant Udgaonkar, Director of Indian Institute of Science Education and Research (IISER), Pune for his assistance with TCSPC facility at National Centre for Biological Sciences (NCBS) Bangalore, India. R.A. acknowledges support of DST, Government of India (EMR/2015/002121), BRNS, Government of India (20150237B02RP00614-BRNS) and DBT, Government of India (BT/PR18927/BCE/8/1376/2016). J.M. acknowledges support of the Department of Atomic Energy, Government of India, under Project Identification No. RTI 4007 and Core Research grants provided by the Department of Science and Technology (DST) of India (CRG/2019/001219).

## ADDITIONAL INFORMATION

Supplementary File.

Contents of file: Supplementary figures S1-S12, supplementary tables S1-S8,.

## Author Contributions

J.S. and M.S. have contributed equally to this work.

## Notes

The authors declare no competing financial interest.

