## Supplementary file for "Phenol Sensing in Nature Modulated via a Conformational Switch Governed by Dynamic Allostery"

[1]Department of Chemistry, Indian Institute of Technology Bombay, Mumbai-400076, India,
[2]Centre for Interdisciplinary Science, Tata Institute of Fundamental Research, Hyderabad-500107, India.
[3]Department of Biochemistry and Molecular Biology, Monash University, Victoria, Australia
[4]Australian Synchrotron, Victoria-3168, Australia,
[5]Department of Biotechnology, Anna University, Chennai, Tamil Nadu 600025, India.
[#] Both authors have equally contributed to the manuscript

**Table of contents**

1. Supplementary figures Pages 2-13
2. Supplementary tables Pages 14-21

### 1. Supplementary figures

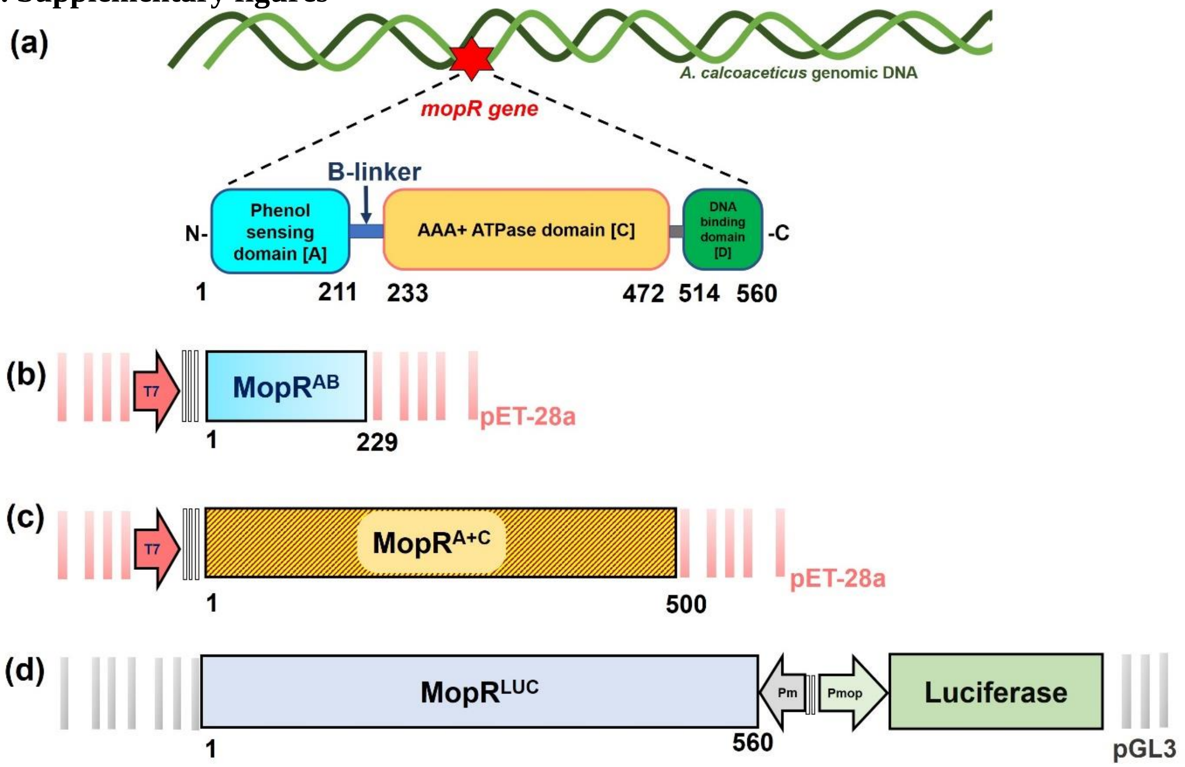


**Figure S1.**
**(a)** Domain organization of MopR from *Acinetobacter calcoaceticus.* Schematic representation of genetic cassette of MopR used **(b)** MopR$^{AB}$ **(c)** MopR$^{A+C}$ **(d)** MopR$^{LUC}$

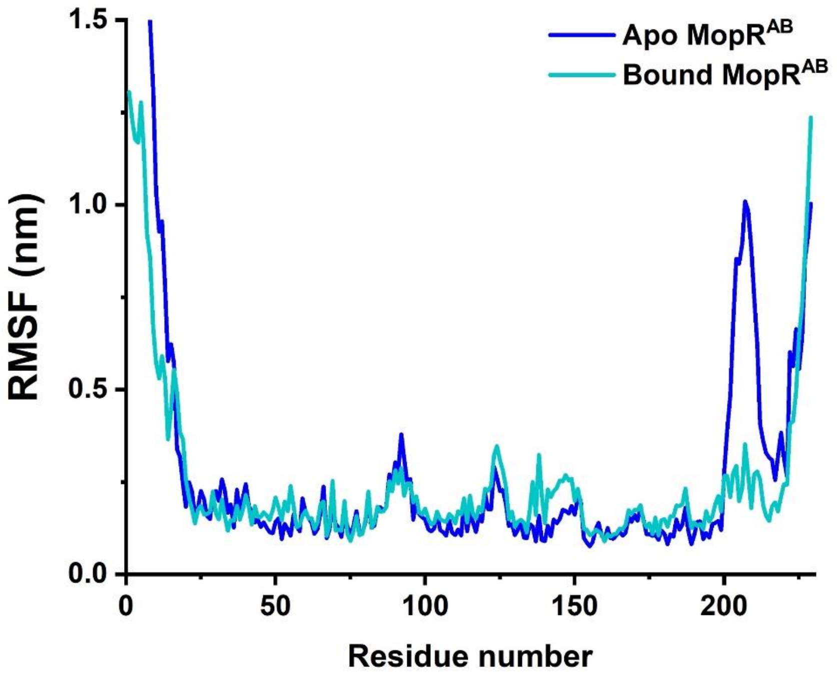


**Figure S2.**
Root Mean Square fluctuations (RMSF) plots of apo and phenol bound states of $MopR^{AB}$

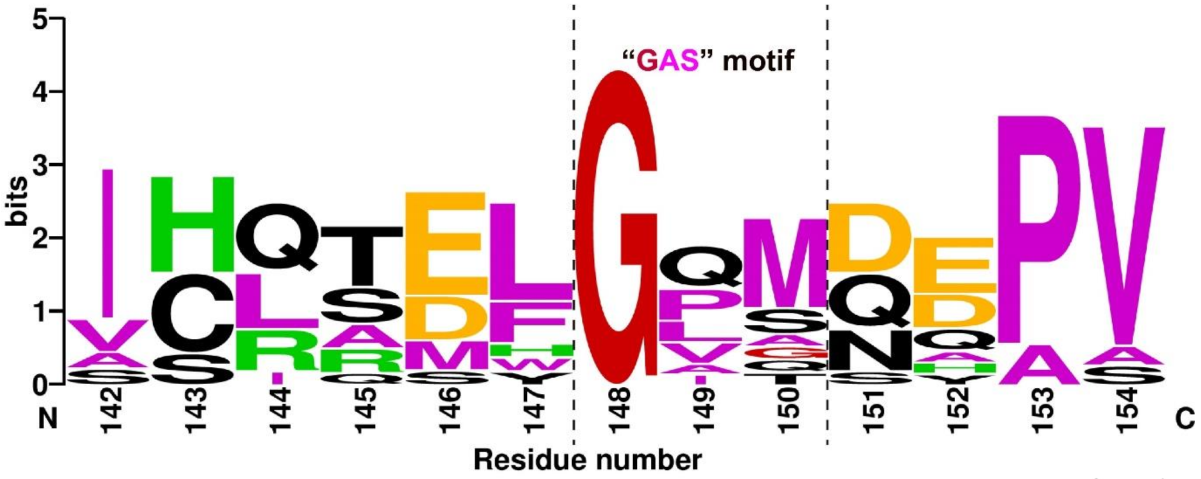


**Figure S3.**
Sequence logo depiction of sequence conservation in aromatic hydrocarbon selective NtrC family proteins (Table S1)

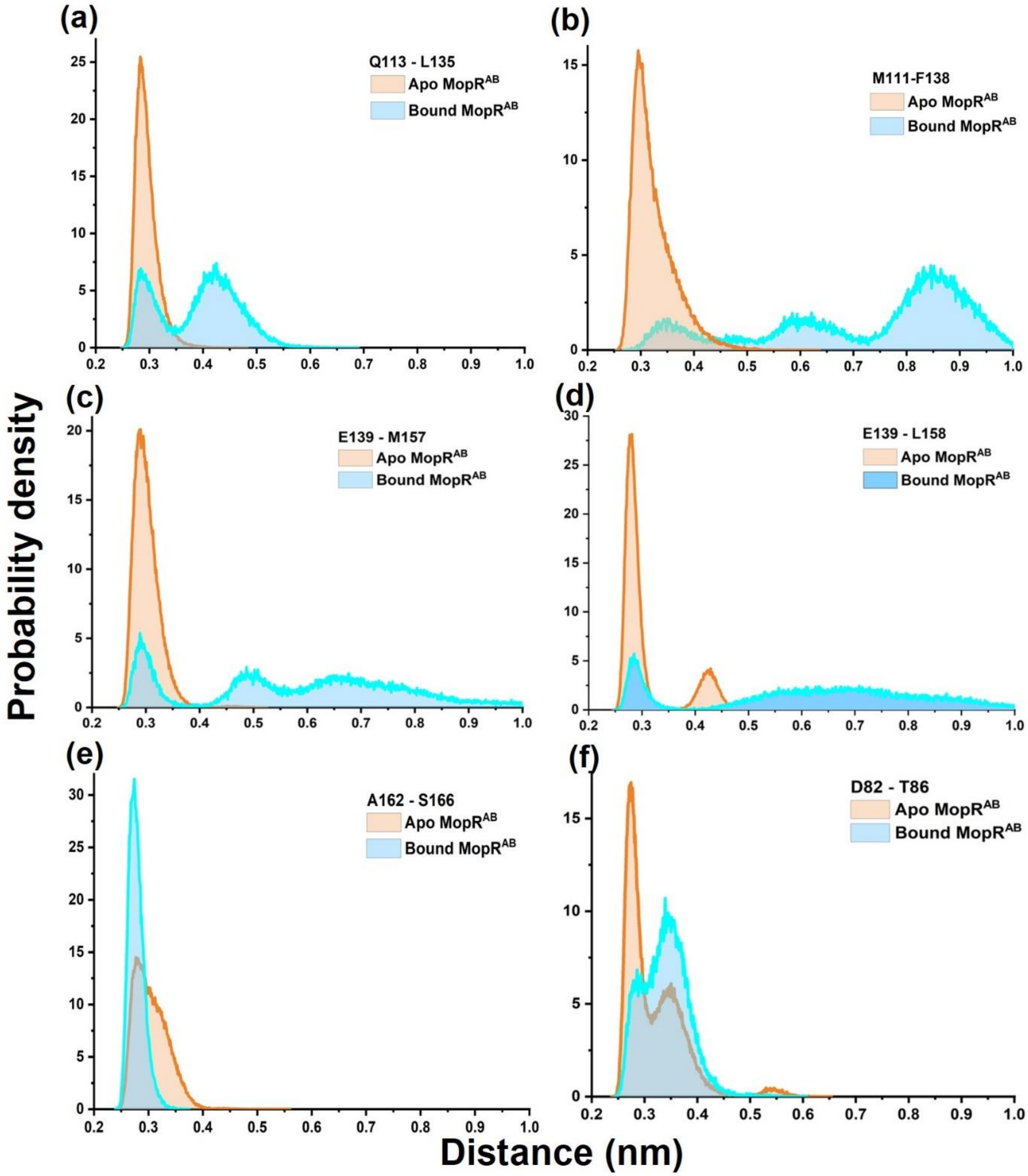


**Figure S4.**
Probability density for the significant inter-residue hydrogen bond distance (nm).

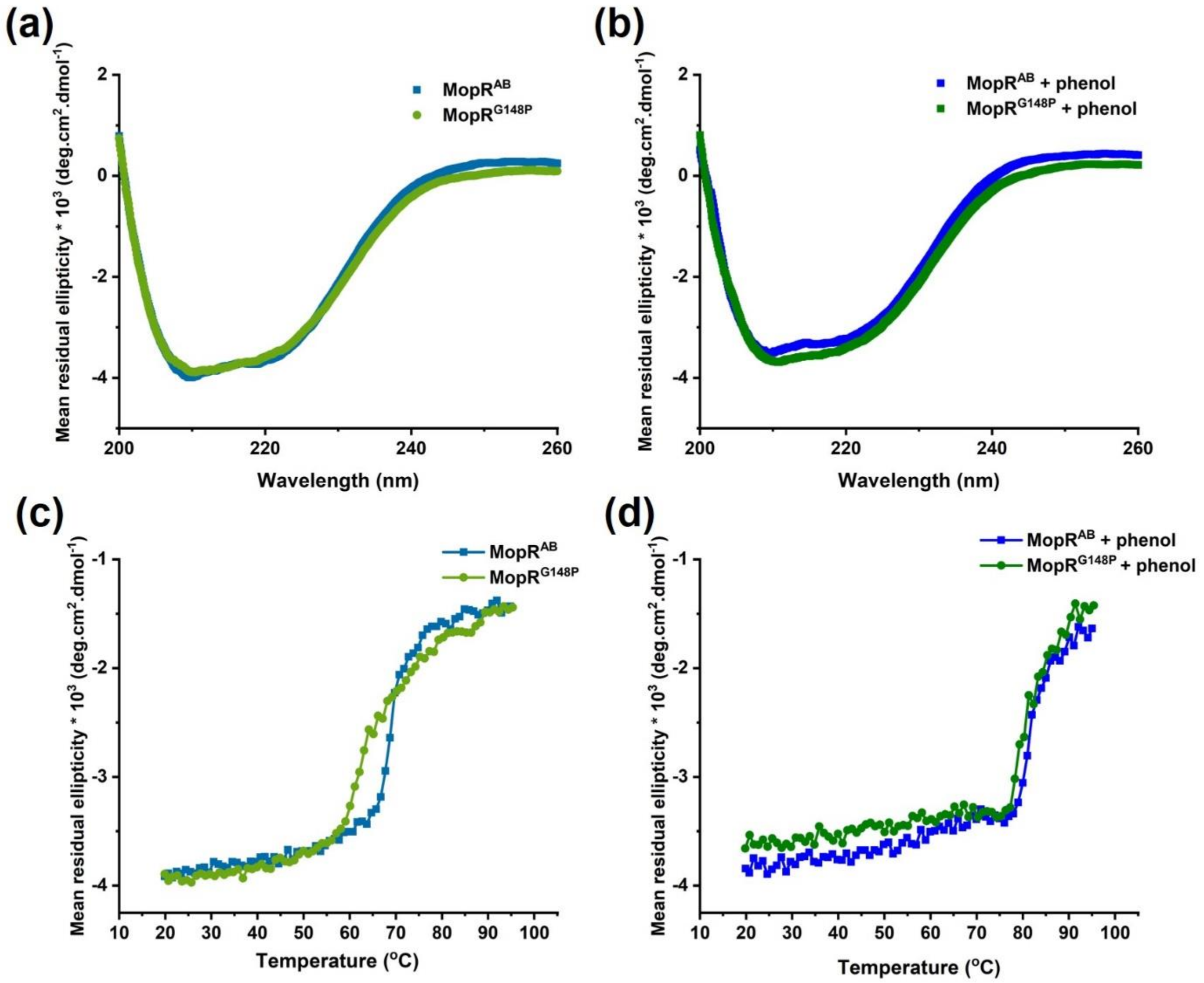


**Figure S5.**
Comparative CD **(a,b)** and CD based Tm spectra **(c,d)** spectra of native MopR$^{AB}$ and MopR$^{G148P}$ with and without phenol.

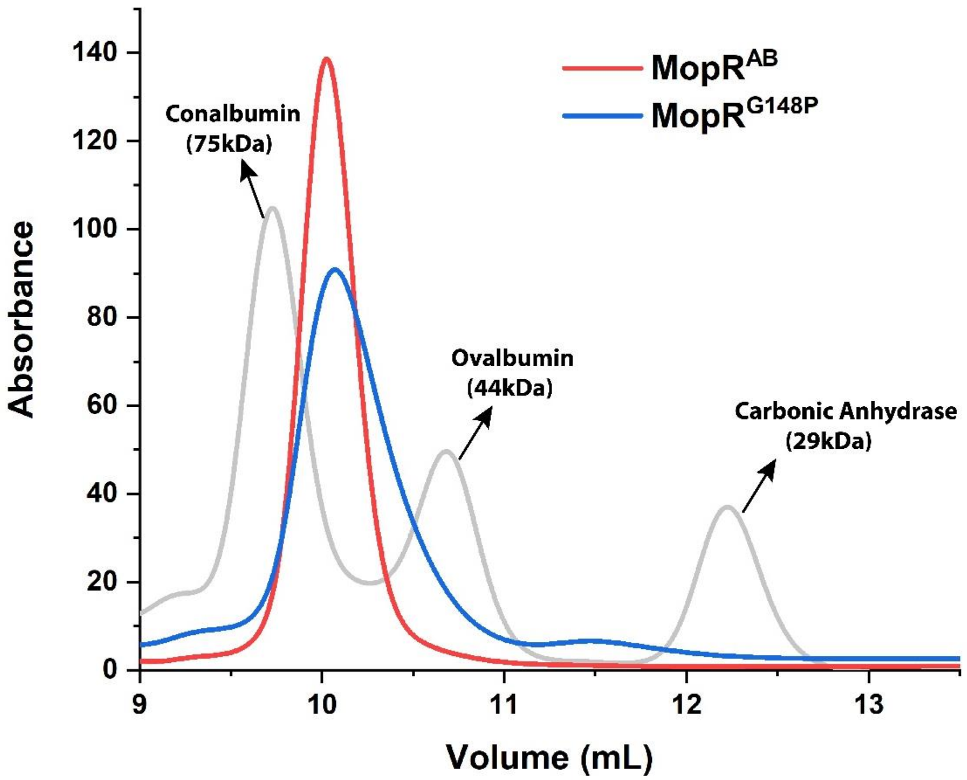


**Figure S6.**
Size Exclusion Chromatography (SEC) profile of $MopR^{AB}$ and $MopR^{G148P}$ using Superdex 75 10/300 GL column. To gauge the molecular weight of MopR, the profile for standards ran on the same column at identical flow rate is depicted in grey.

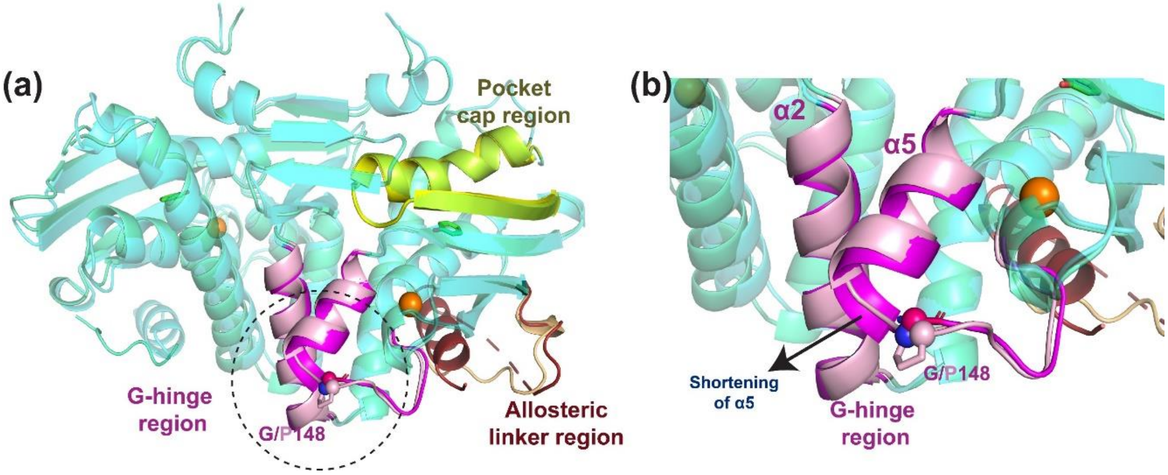


**Figure S7.**
**(a)** Structural superposition of phenol bound states of MopR$^{AB}$ (in palecyan) and MopR$^{G148P}$ (in aquagreen) highlighting the significant regions. **(b)** Magnified view depicting rearrangement in the G-hinge region.

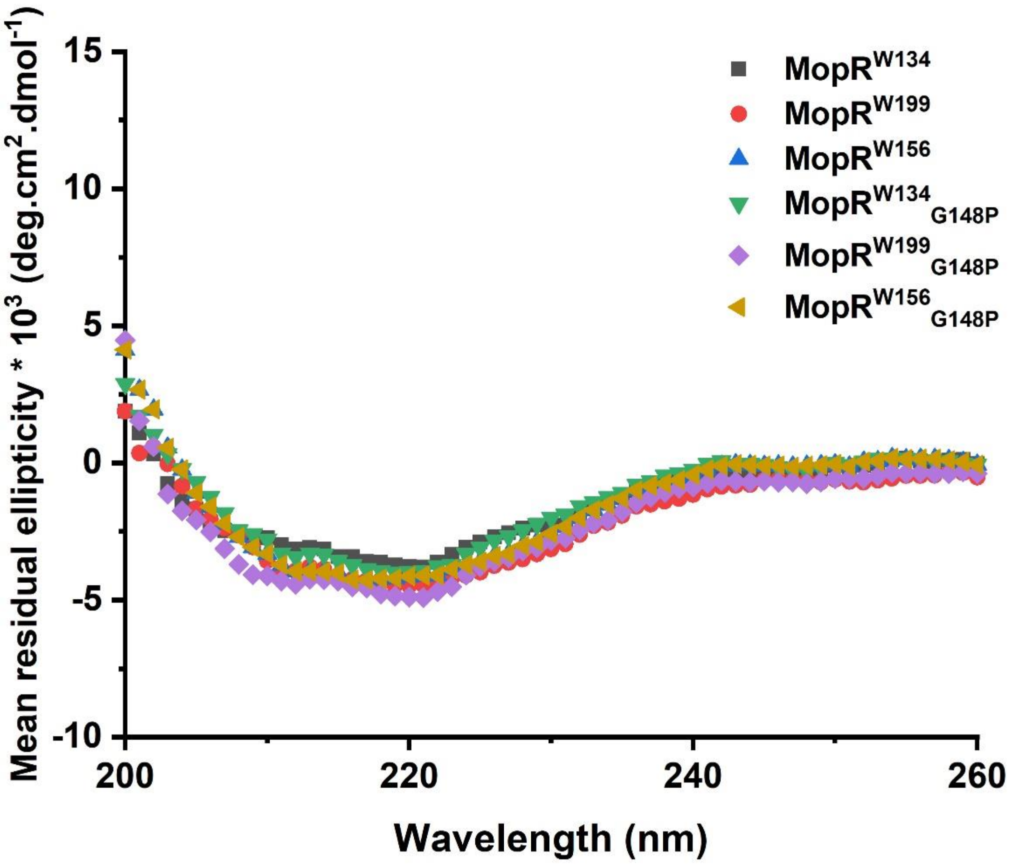


**Figure S8.**
CD spectrum of various single tryptophan mutants.

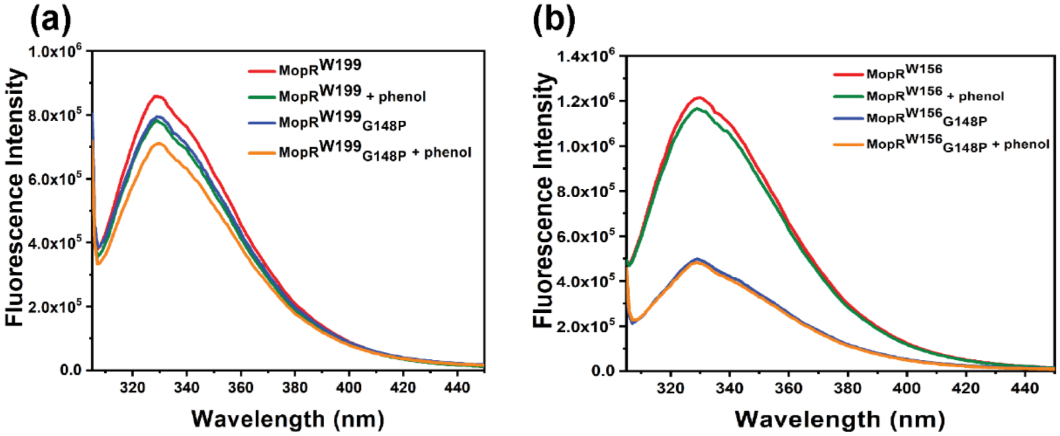


**Figure S9.**
Steady state fluorescence studies for various single tryptophan mutants. **(a, b)** Emission spectra of **(a)** MopR$^{W199}$ and MopR$^{W199}{}_{G148P}$ **(b)** MopR$^{W156}$ and MopR$^{W156}{}_{G148P}$, in presence and absence of phenol.

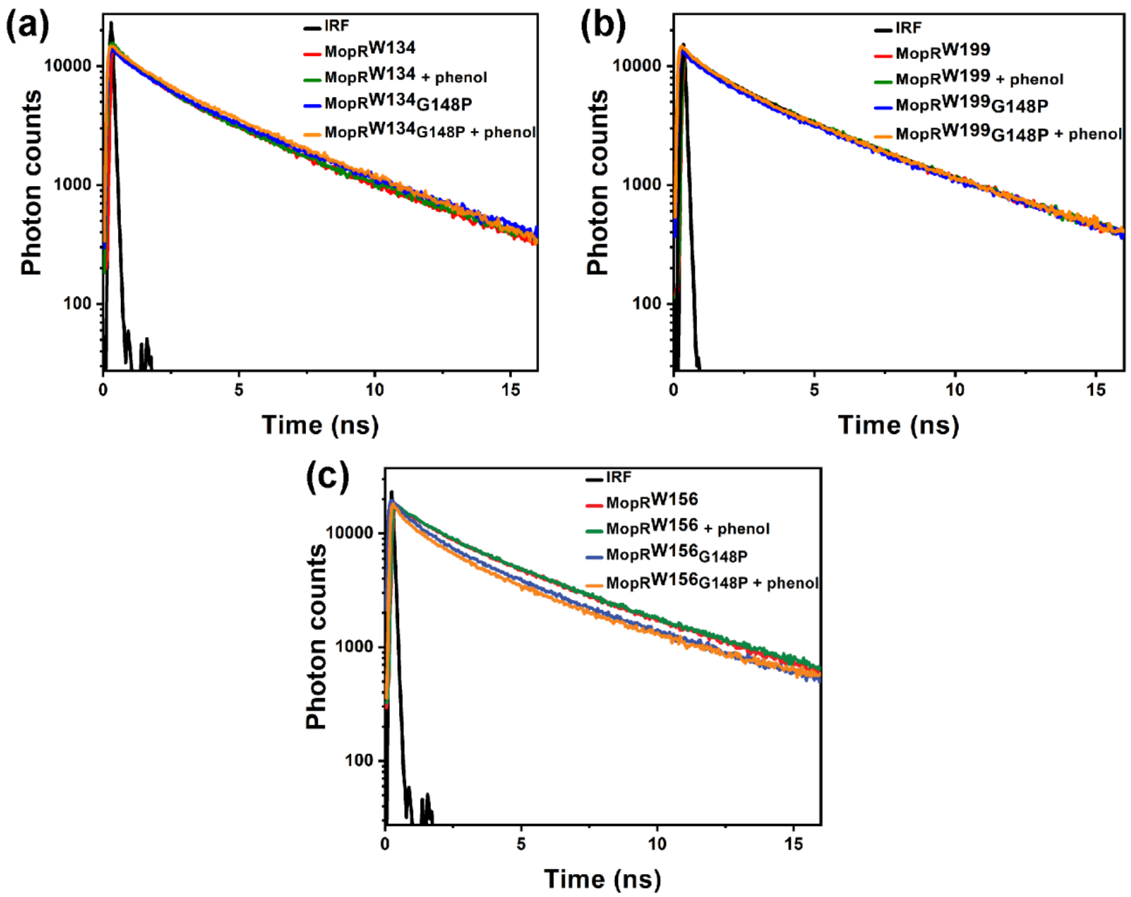


**Figure S10.**
Intensity decay curves of **(a)** $MopR^{W134}$ and $MopR^{W134}{}_{G148P}$ and **(b)** $MopR^{W199}$ and $MopR^{W199}{}_{G148P}$ **(c)** $MopR^{W156}$ and $MopR^{W156}{}_{G148P}$ in presence and absence of phenol**.**

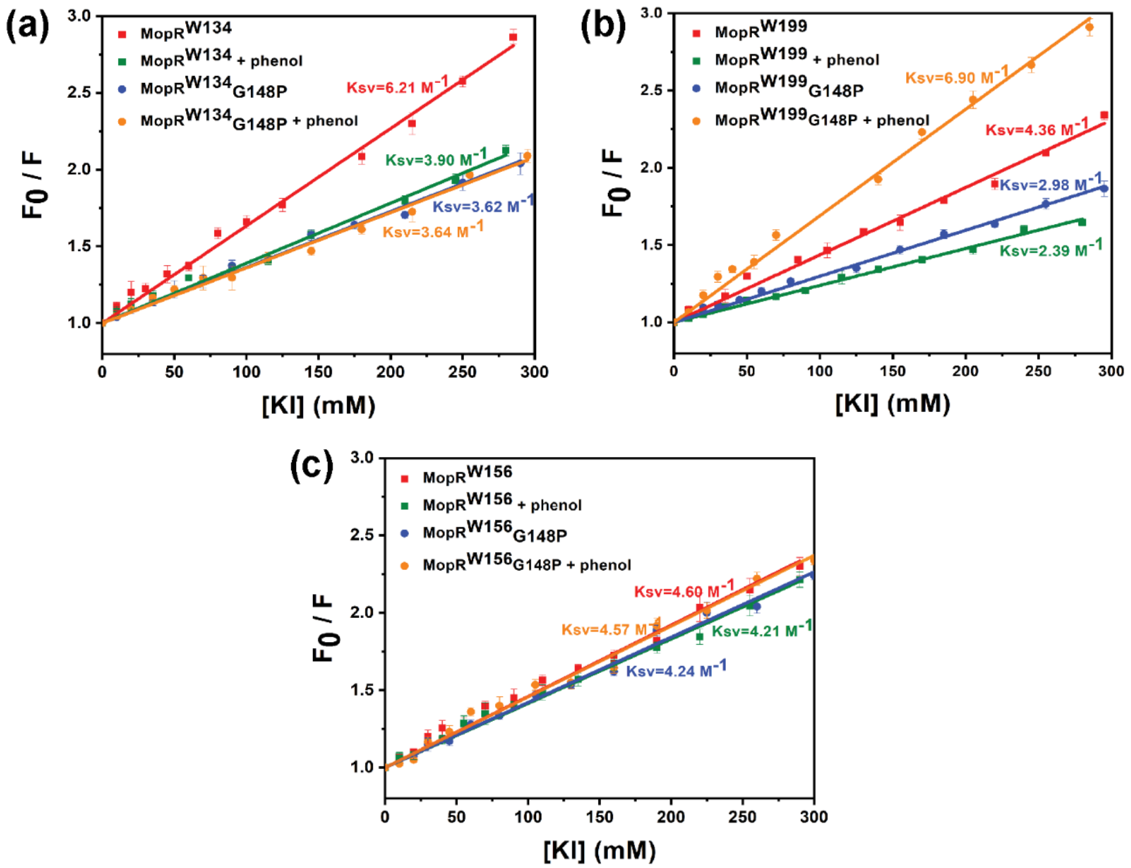


**Figure S11.**
Stern-Volmer plots for **(a)** MopR$^{W134}$ and MopR$^{W134}$$_{G148P}$ **(b)** MopR$^{W199}$ and MopR$^{W199}$$_{G148P}$ and **(c)** MopR$^{W156}$ and MopR$^{W156}$$_{G148P}$, in presence and absence of phenol.

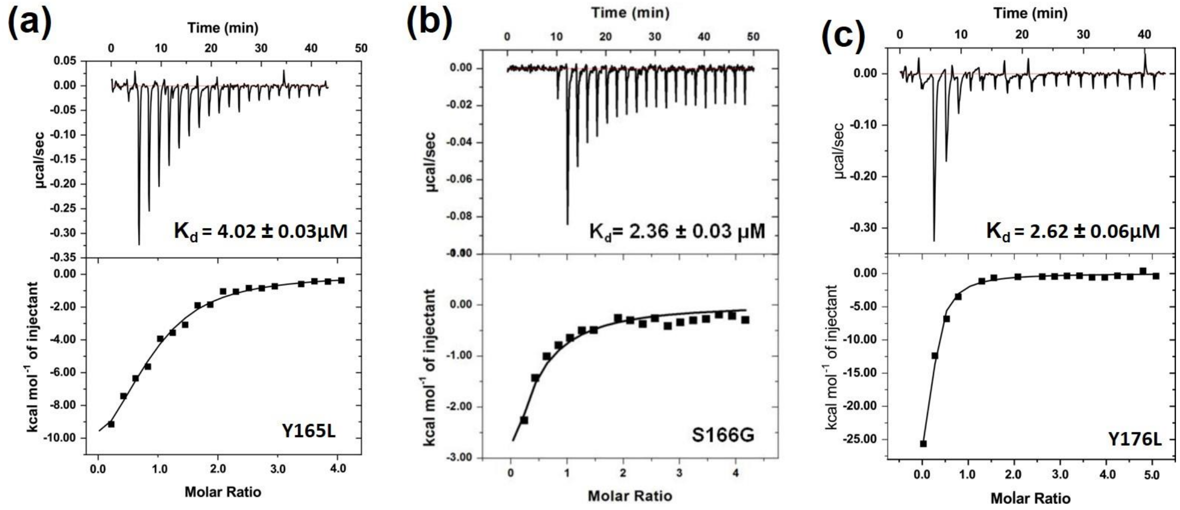


**Figure S12.**
ITC studies to determine affinity of phenol towards MopR$^{AB}$ mutants **(a)** Y165L, **(b)** S166G and **(c)** Y176L.

### 2. Supplementary tables.

**Table S1.**
Database details of the NtrC family proteins used for the bioinformatic studies.

| Protein | Microorgarism | Primary Effector | Accession number |
|---|---|---|---|
| **MopR** | *Acinetobacter calcoaceticus NCIB8250* | Phenol | CAA93242.1 |
| **PoxR** | *Ralstoni eutropha E2* | Phenol | AAC32451 |
| **CapR** | *Pseudomonas putida KCTC 1452* | Phenol | AAP46187 |
| **AphR** | *Comamonas testosterone TA441* | Phenol | BAA34177 |
| **PheR** | *Pseudomonas putida BH* | Phenol | BAA09883 |
| **PhcR** | *Comamonas testosterone R5* | Phenol | BAA87867 |
| **PhlR** | *Pseudomonas putida H (pPGH1)* | Catechol | CAA56747 |
| **DmpR** | *Pseudomonas sp. CF600(pVI150)* | 2,3- dimethylphenol | A47078 |
| **XylR** | *Pseudomonas putida* mt-2 | BTEX | AAA26028 |
| **TbuT** | *Ralstonia pickettii PKO1* | BTEX | AAC44567 |
| **BhpR** | *Novosphingobium aromaticivorans* | Biphenyl | AAD03979 |
| **HbpR** | *Pseudomonas nitroreducens* | 2-hydroxybiphenyl | O06645 |
| **PhnR** | *Burkholderia sartisoli* | Naphthalene/ Phenanthrene | Q9ZHH9 |
| **TmbR** | *Pseudomonas putida TMB* | Trimethylbenzene | U41301 |

**Table S2.**
Hydrogen bond propensities in the vicinity of "pocket cap region" as measured over different system states.

| **Donor** | **Acceptor** | **Apo MopR$^{AB}$** | **Bound MopR$^{AB}$** |
|---|---|---|---|
| TYR165 | ALA98 | 0.42 | 0.94 |
| THR86 | TYR165 | 0.34 | 0.67 |
| THR86 | ASP82 | 0.54 | 0.25 |
| SER166 | ALA162 | 0.47 | 0.91 |
| HIS106 | VAL112 | 0.00 | 0.64 |
| GLN113 | LEU135 | 0.96 | 0.31 |
| PHE138 | MET111 | 0.60 | 0.04 |
| GLN177 | TYR176 | 0.02 | 0.54 |
| GLN177 | ILE218 | 0.11 | 0.00 |
| GLN177 | GLN222 | 0.24 | 0.01 |

**Table S3.**
Hydrogen bond propensities in the vicinity of “G-hinge region” as measured over different system states

| **Donor** | **Acceptor** | **Apo MopR$^{AB}$** | **Bound MopR$^{AB}$** |
|---|---|---|---|
| ARG55 | GLU139 | 0.62 | 0.21 |
| ARG55 | GLU139 | 0.11 | 0.01 |
| ARG55 | HIS143 | 0.21 | 0.06 |
| HIS143 | GLY148 | 0.45 | 0.11 |
| TRP156 | GLU139 | 0.74 | 0.17 |
| TRP156 | GLU139 | 0.42 | 0.10 |
| TRP156 | ALA154 | 0.00 | 0.26 |
| MET157 | GLU139 | 0.20 | 0.03 |
| MET157 | GLU139 | 0.80 | 0.20 |
| LEU158 | GLU139 | 0.82 | 0.20 |
| LEU159 | CYS155 | 0.00 | 0.35 |
| GLU141 | SER137 | 0.81 | 0.51 |

**Table S4.**
Hydrogen bond propensities in the vicinity of “allosteric linker region” as measured over different system states.

| **Donor** | **Acceptor** | **Apo MopR$^{AB}$** | **Bound MopR$^{AB}$** |
|---|---|---|---|
| TRP199 | ILE175 | 0.63 | 0.00 |
| TRP199 | LYS194 | 0.24 | 0.00 |
| ILE206 | ALA202 | 0.00 | 0.85 |
| PHE208 | GLU204 | 0.00 | 0.72 |
| MET209 | LEU205 | 0.00 | 0.95 |
| SER210 | ILE206 | 0.00 | 0.48 |
| SER210 | ILE206 | 0.00 | 0.50 |

**Table S5.**
Binding pocket volume calculation as measured over different system states using MDpocket.

| SYSTEM STATE | POCKET VOLUME ($A^3$) |
|---|---|
| **Apo MopR$^{AB}$** | 158.3 ± 1.1 |
| **Bound MopR$^{AB}$** | 221.9 ± 0.8 |

**Table S6.**
Stacking interactions in binding domain in bound MopR$^{AB}$ state.

| | PHE99 | PHE128 | PHE132 | TYR161 | TYR165 | TYR176 | HIS106 | TRP134 | Phenol |
|---|---|---|---|---|---|---|---|---|---|
| **PHE99** | - | | | | | | | | |
| **PHE128** | 1.00 | - | | | | | | | |
| **PHE132** | 0.86 | 0.00 | - | | | | | | |
| **TYR161** | 0.00 | 0.00 | 0.00 | - | | | | | |
| **TYR165** | 0.10 | 0.00 | 0.00 | 0.99 | - | | | | |
| **TYR176** | 0.02 | 0.79 | 0.94 | 0.00 | 0.00 | - | | | |
| **HIS106** | 0.00 | 0.00 | 0.00 | 1.00 | 0.00 | 0.00 | - | | |
| **TRP134** | 0.00 | 0.00 | 0.86 | 0.00 | 0.00 | 0.00 | 0.67 | - | |
| **Phenol** | 1.00 | 0.00 | 1.00 | 0.74 | 1.00 | 0.69 | 1.00 | 0.16 | - |

**Table S7.**
Thermodynamic parameters of binding of $MopR^{AB}$ and $MopR^{G148P}$ mutant with phenol using ITC studies.

| **Construct** | **ΔH (kcal/mol)** | **TΔS (kcal/mol)** | **ΔG (kcal/mol)** | **$K_d$ (μM)** | **$N^a$** | **c-value$^b$** |
|---|---|---|---|---|---|---|
| $MopR^{AB}$ | -17.6 ± 0.13 | -8.64 ± 0.18 | -8.98 ± 0.06 | 0.46 ± 0.06 | 1.11 | 86.9 |
| $MopR^{G148P}$ | -14.7 ± 0.32 | -5.36 ± 0.30 | -9.21 ± 0.12 | 0.07 ± 0.02 | 1.02 | 142.8 |

$^a$N = Number of sites
$^b$c-value = n [Protein]/ $K_d$

**Table S8.**

Lifetimes and their amplitudes of various single tryptophan mutants of $MopR^{AB}$ and $MopR^{G148P}$.

| Protein | $\tau_1$(ns) | $\alpha_1$ | $\tau_2$(ns) | $\alpha_2$ | $\tau_3$(ns) | $\alpha_3$ | $\tau_m$(ns) | $\chi2$ |
|---|---|---|---|---|---|---|---|---|
| **$MopR^{W199}$** | 0.48 (±0.01) | 0.15 (±0.04) | 1.71 (±0.08) | 0.29 (±0.03) | 5.00 (±0.02) | 0.53 (±0.01) | 3.28 (±0.07) | 1.07 |
| **$MopR^{W199}$+phenol** | 0.41 (±0.08) | 0.17 (±0.01) | 1.78 (±0.01) | 0.34 (±0.02) | 5.06 (±0.03) | 0.49 (±0.02) | 3.15 (±0.06) | 1.10 |
| **$MopR^{W156}$** | 0.87 (±0.11) | 0.14 (±0.02) | 1.91 (±0.09) | 0.31 (±0.03) | 5.19 (±0.02) | 0.56 (±0.01) | 3.59 (±0.05) | 1.12 |
| **$MopR^{W156}$+phenol** | 0.75 (±0.01) | 0.15 (±0.10) | 1.92 (±0.01) | 0.32 (±0.01) | 5.19 (±0.04) | 0.53 (±0.02) | 3.47 (±0.08) | 1.09 |
| **$MopR^{W134}$** | 0.34 (±0.07) | 0.15 (±0.02) | 1.38 (±0.03) | 0.41 (±0.01) | 4.70 (±0.04) | 0.44 (±0.01) | 2.70 (±0.02) | 1.18 |
| **$MopR^{W134}$+phenol** | 0.47 (±0.06) | 0.20 (±0.06) | 1.52 (±0.14) | 0.41 (±0.05) | 4.89 (±0.01) | 0.39 (±0.01) | 2.62 (±0.02) | 1.05 |
| **$MopR^{W199}_{G148P}$** | 0.17 (±0.03) | 0.18 (±0.04) | 1.10 (±0.03) | 0.29 (±0.02) | 4.44 (±0.09) | 0.52 (±0.01) | 2.68 (±0.02) | 1.16 |
| **$MopR^{W199}_{G148P}$+phenol** | 0.13 (±0.04) | 0.24 (±0.03) | 1.08 (±0.06) | 0.29 (±0.05) | 4.40 (±0.10) | 0.47 (±0.02) | 2.41 (±0.11) | 1.15 |
| **$MopR^{W156}_{G148P}$** | 0.11 (±0.01) | 0.31 (±0.02) | 1.08 (±0.04) | 0.30 (±0.03) | 4.28 (±0.02) | 0.39 (±0.01) | 2.03 (±0.05) | 1.02 |
| **$MopR^{W156}_{G148P}$+phenol** | 0.15 (±0.02) | 0.30 (±0.02) | 1.14 (±0.04) | 0.30 (±0.01) | 4.34 (±0.06) | 0.39 (±0.04) | 2.09 (±0.06) | 1.12 |
| **$MopR^{W134}_{G148P}$** | 0.20 (±0.04) | 0.17 (±0.03) | 1.24 (±0.06) | 0.29 (±0.02) | 4.42 (±0.05) | 0.54 (±0.01) | 2.79 (±0.04) | 1.02 |
| **$MopR^{W134}_{G148P}$+phenol** | 0.18 (±0.02) | 0.16 (±0.03) | 1.28 (±0.05) | 0.30 (±0.01) | 4.48 (±0.01) | 0.53 (±0.01) | 2.80 (±0.01) | 1.02 |

All measurements were carried out in triplicates and mean ± SE (standard error) have been reported
